# ICOS signaling limits regulatory T cell accumulation and function in visceral adipose tissue

**DOI:** 10.1101/2020.06.01.128504

**Authors:** Kristen L. Mittelsteadt, Erika T. Hayes, Daniel J. Campbell

## Abstract

A unique population of Foxp3^+^ regulatory T cells (T_R_) resides in visceral adipose tissue (VAT) that regulates adipose inflammation and helps preserve insulin sensitivity. The costimulatory molecule ICOS is highly expressed on effector (e)T_R_ that migrate to nonlymphoid tissues, and contributes to their maintenance and function in models of autoimmunity. In this study, we report an unexpected cell-intrinsic role for ICOS expression and downstream PI3K signaling in limiting the abundance, VAT-associated phenotype, and function of T_R_ specifically in VAT. *Icos*^*−/−*^ mice and mice expressing a knock-in form of ICOS that cannot activate PI3K had increased VAT-T_R_ abundance and elevated expression of canonical VAT-T_R_ markers. Loss of ICOS signaling facilitated enhanced accumulation of T_R_ to VAT associated with elevated CCR3 expression, and resulted in reduced adipose inflammation and heightened insulin sensitivity in the context of high-fat diet. Thus, we have uncovered a new and surprising molecular pathway that regulates VAT-T_R_ accumulation and function.

**Summary:** The authors demonstrate that loss of ICOS-dependent PI3-kinase signaling supports visceral adipose tissue (VAT) T_R_ abundance and function, and this correlates with reduced adipose inflammation and improved insulin sensitivity after high-fat diet. This work highlights a new molecular pathway that regulates VAT-T_R_ accumulation and activity.

## Introduction

CD4^+^Foxp3^+^ regulatory T cells (T_R_) are critical for maintaining immune tolerance and for the resolution of ongoing inflammation after infection (Dominguez-Villar and Hafler, 2018; Rivas and Chatila, 2016; Smigiel et al., 2014a). Specialized subsets of tissue-specific T_R_ also function in tissue repair and homeostasis. For example, Areg^+^ T_R_ expand in skeletal muscle and the lungs in response to injury and are required for proper tissue repair (Arpaia et al., 2015; Burzyn et al., 2013), whereas T_R_ within hair follicles and skin are essential for hair generation and inhibiting fibrosis (Ali et al., 2017; Kalekar et al., 2019). T_R_ are also found in the visceral adipose tissue (VAT) of both mice and humans, where they regulate adipose inflammation and preserve metabolic homeostasis (Deiuliis et al., 2011; Eller et al., 2011; Feuerer et al., 2009; Ilan et al., 2010). As such, T_R_ ablation in diphtheria toxin-treated *Foxp3*^DTR^ mice results in increased inflammatory mediators in VAT and reduced insulin sensitivity, whereas augmentation of T_R_ by administration of IL-2 immune complexes or IL-33 results in improved metabolic readouts in mice on high-fat diet (HFD) (Feuerer et al., 2009; Han et al., 2015; Vasanthakumar et al., 2015).

Given that T_R_ modulate diverse responses in different anatomical and inflammatory settings, it is not surprising that they exhibit considerable phenotypic and functional heterogeneity. Expression of CD62L and CD44 broadly divides T_R_ into distinct subsets that are enriched in lymphoid and nonlymphoid tissues (Campbell, 2015; Smigiel et al., 2014b). We term these populations central T_R_ (cT_R_) and effector T_R_ (eT_R_), respectively. Differential activation of PI3K, unique epigenetic landscapes, and distinct transcriptional programs underlie the diversification of T_R_, and selective expression of chemokine receptors and cell adhesion molecules give subsets of T_R_ access to specific tissues (Delacher et al., 2017; DiSpirito et al., 2018; Luo et al., 2016). The proper distribution of T_R_ across tissues is crucial for maintaining immune tolerance and tissue homeostasis (Sather et al., 2007; Yamaguchi et al., 2011). Hence, understanding the signals that control the development, maintenance, and function of tissue-specific T_R_ is vital to fully harnessing their therapeutic potential.

T_R_ occupancy in different tissues is met with unique homeostatic maintenance requirements. Generally, IL-2 signaling maintains cT_R_ within T cell zones of secondary lymphoid tissues by driving pro-survival signals, whereas maintenance of eT_R_ in nonlymphoid tissues can be IL-2 independent and instead relies on TCR and costimulatory molecule engagement (Levine et al., 2014; Smigiel et al., 2014b; Tang et al., 2003). ICOS is expressed on highly suppressive T_R_ and can control T_R_ abundance by inhibiting apoptosis and stimulating proliferation (Burmeister et al., 2008; Redpath et al., 2013). IL-10 is essential for control of local immune responses, and is primarily expressed by Blimp-1^+^ICOS^+^ eT_R_ (Cretney et al., 2011). ICOS costimulation *in vitro* super-induces IL-10 expression (Arimura et al., 2002; Hutloff et al., 1999), and this depends on its ability to activate phosphoinositide 3-kinase (PI3K) (Feito et al., 2003; Okamoto et al., 2003). Although mice deficient in ICOS or ICOS ligand do not develop spontaneous inflammatory or autoimmune disease, genetic or antibody-mediated blockade of ICOS signaling promotes inflammation in mouse models of autoimmunity and infection as a result of reduced T_R_ abundance and/or IL-10 production (Busse et al., 2012; Dong et al., 2001; Herman et al., 2004; Kohyama et al., 2004; Kornete et al., 2012; Landuyt et al., 2019; Miyamoto et al., 2005; Redpath et al., 2013).

ICOS potently activates PI3K and downstream AKT via a unique YMFM motif in its cytoplasmic tail (Arimura et al., 2002; Fos et al., 2008). AKT can act on several different substrates, including the transcription factor Foxo1, to modulate transcription of genes involved in proliferation, survival, metabolic reprogramming, and migration/tissue tropism (So and Fruman, 2012). Indeed, inactivation of Foxo1 by AKT is essential for the differentiation of eT_R_ and their migration to nonlymphoid tissues (Luo et al., 2016). <u>The mammalian target of rapamycin (mTOR) pathway is also activated by PI3K signaling, and regulates Treg cell metabolism and differentiation</u>. Thus, engagement of ICOS may facilitate the development and/or maintenance of eT_R_ via multiple PI3K-dependent signaling pathways.

Accumulation of T_R_ in VAT is dependent on several signals. VAT-T_R_ consist of clonally expanded populations, suggesting recognition of one or more adipose tissue antigens promotes their tissue residence (Feuerer et al., 2009; Kolodin et al., 2015). Additionally, mice lacking expression of IL-33 or its receptor ST2 are deficient in VAT-T_R_, whereas injection of exogenous IL-33 results in increased VAT-T_R_ abundance with little impact on lymphoid T_R_ (Han et al., 2015; Vasanthakumar et al., 2015). Recent work also identified a cell-intrinsic role for CCR2 in the ability of donor T_R_ to repopulate VAT (Vasanthakumar et al., 2020). Finally, T_R_ expression of PPARγ, the master regulator of adipocyte differentiation, supports VAT-T_R_ phenotype and accumulation and drives expression of factors important for lipid metabolism (Cipolletta et al., 2012; Cipolletta et al., 2015). However, despite having an eT_R_ phenotype, the role of costimulatory molecule signaling in VAT-T_R_ is poorly understood.

In this study, we demonstrate an unexpected role for cell-intrinsic ICOS-dependent PI3K signaling in restricting VAT-T_R_ accumulation and function. Moreover, we implicate the CCL11/24-CCR3 axis as an additional factor capable of driving recruitment of T_R_ to VAT, which is enhanced in the absence of ICOS signaling. These surprising findings challenge the current model regarding signals that support diverse T_R_ subsets, and highlight the cell- and tissue-specific effects of ICOS expression and signaling on T_R_ development, accumulation, and function.

## Results

### Mice lacking ICOS signaling have reduced T_R_ in lymphoid tissues and altered expression of PI3K targets

To determine how ICOS signaling impacts T_R_ abundance in different tissues, we crossed *Foxp3*^*mRFP*^ mice to *Icos*^*−/−*^ (KO) mice and to *Icos*^*Y181F*^ (YF) mice, which carry a tyrosine-to-phenylalanine knock-in mutation in the YMFM motif in the cytoplasmic tail of ICOS, thereby specifically abolishing ICOS-mediated PI3K activation (Gigoux et al., 2009), to *Foxp3*^*mRFP*^ mice (Wan and Flavell, 2005). In line with published data, we found a ~30% reduction in the frequency and number of T_R_ in the spleens of KO compared to WT *Foxp3*^*mRFP*^ mice (Burmeister et al., 2008; Landuyt et al., 2019) **[Fig. 1A]**. This was even more pronounced in older animals, as T_R_ accumulate with age (Nishioka et al., 2006) **[Fig. 1B]**. Despite normal ICOS expression on the cell surface **[Fig. S1A]** and intact MAPK signaling (Gigoux et al., 2009), we saw a similar reduction in splenic T_R_ abundance in YF mice, indicating that activation of PI3K signaling is the critical pathway by which ICOS regulates T_R_ abundance **[Fig. 1A, 1B]**. ICOS is expressed on cT_R_, but its expression is further upregulated on eT_R_ **[Fig. S1B]**. The loss of T_R_ we observed in YF and KO spleens was associated with a disproportionate reduction in the frequency and number of eT_R_, whereas cT_R_ abundance was unchanged, consistent with results using antibody blockade of ICOS signaling (Smigiel et al., 2014b) **[Fig. 1C]**. Loss of T_R_ was also observed in the peripheral lymph nodes (<u>pLN</u>) of YF and KO mice, indicating that ICOS signaling is important for eT_R_ maintenance in multiple secondary lymphoid organs **[Fig. S1C]**. Both YF and KO T_R_ in the spleen had elevated expression of the lymphoid homing molecules CD62L and CCR7 which are downregulated upon PI3K activation via phosphorylation and sequestration of Foxo1 (Kerdiles et al., 2009) **[Fig. 1D]**. This reflects both an increased frequency of splenic cT_R_ as well as increased CD62L and CCR7 expression by gated CD44^hi^ T_R_ in YF and KO mice. When examined directly *ex vivo*, YF and KO splenic T_R_ also exhibited reduced phosphorylation of the mTORC1 target, ribosomal protein S6, consistent with diminished activation of PI3K **[Fig. 1E]**. Thus, the absence of ICOS-mediated PI3K activation drives the altered lymphoid T_R_ frequency and phenotype we see in YF and KO mice.

**Figure 1:**
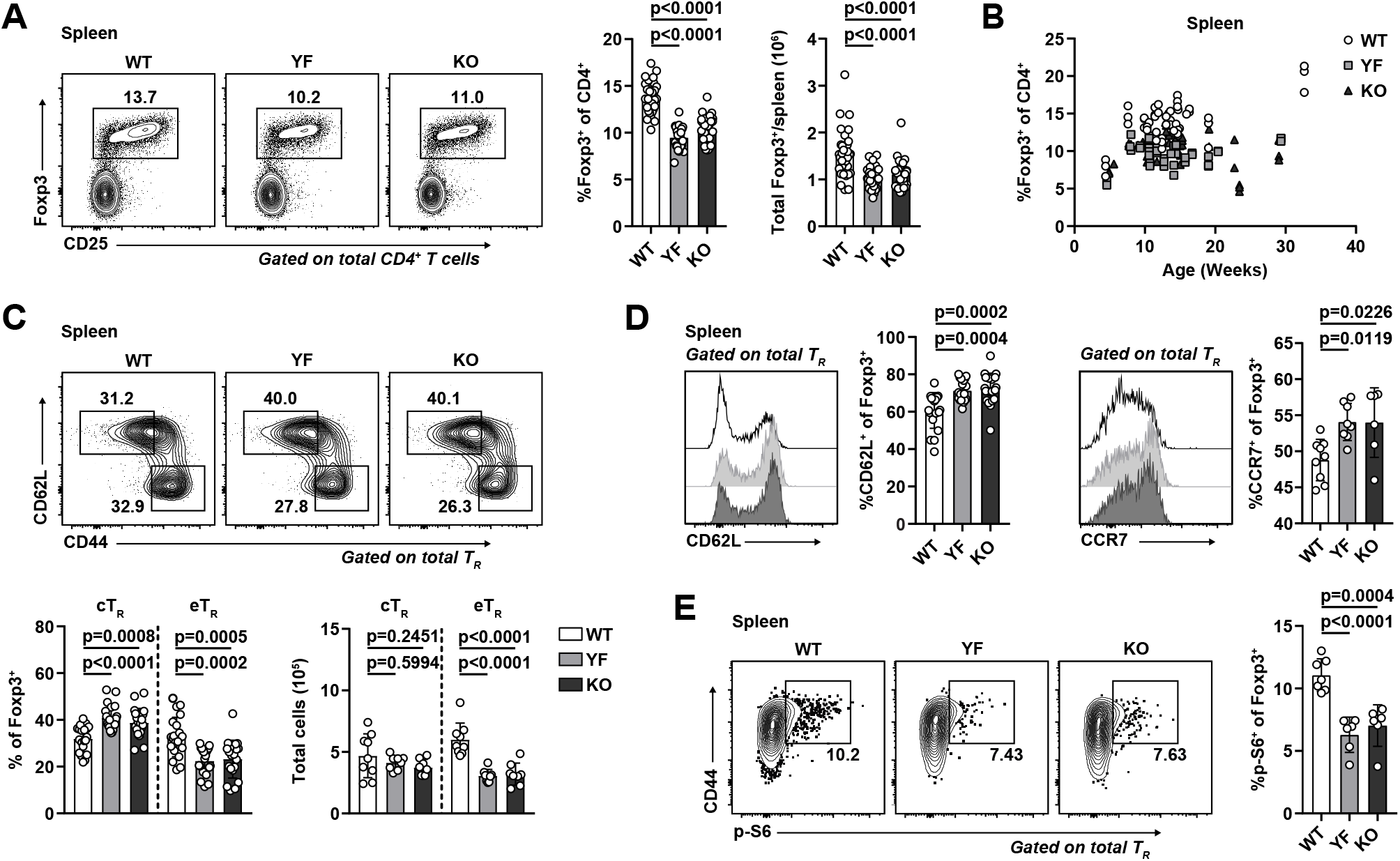
Absence of ICOS signaling results in loss of lymphoid T_R_ and altered expression of PI3K targets. **(A)** Representative flow cytometry plots and quantification of splenic T_R_ (*n* = 3-6 per group from 9 independent experiments). **(B)** Splenic T_R_ frequencies at indicated ages (*n* = 3-5 per group from 16 independent experiments). **(C)** Representative flow cytometry plots and quantification of splenic cT_R_ (CD44^lo^CD62L^hi^) and eT_R_ (CD44^hi^CD62L^lo^) (*n* = 3-5 per group from 7 independent experiments). **(D)** Expression of CD62L and CCR7 in splenic T_R_ (CD62L: *n* = 3-5 per group from 5 independent experiments. CCR7: *n* = 3-5 per group from 2 independent experiments). **(E)** S6 phosphorylation measured directly *ex vivo* in splenic T_R_ by flow cytometry. (*n* = 3-4 per group from 2 independent experiments). Mice were age-matched within independent experiments and pooled data are from experiments using mice aged 8-16 weeks, except in **(B)** where age is indicated. Statistical significance was determined using one-way ANOVA with Tukey’s post-test. All graphical data are presented as mean values ± SD.

### T_R_ are enriched in VAT in the absence of ICOS signaling

As eT_R_ are highly enriched in nonlymphoid sites (Lee et al., 2007; Smigiel et al., 2014b), we next assessed whether maintenance of T_R_ in peripheral tissues was diminished in YF and KO mice. Using intravascular labeling to identify tissue-localized T_R_ (Anderson et al., 2014), we observed a reduction in the frequency and number of T_R_ in the skin and decrease in the absolute number of T_R_ in the lungs of YF and KO mice [**Fig. 2A, 2B]**. However, T_R_ frequency and total number were actually elevated in the VAT of both strains **[Fig. 2C]**. This was specific to VAT, as frequency and total number of T_R_ in subcutaneous adipose tissue (SQAT) remained similar between genotypes **[Fig. 2D]**. Male and female YF and KO mice demonstrated increased VAT-T_R_ abundance compared to WT mice of the same sex, however the frequency and number of VAT-T_R_ in females was significantly less than that observed in males **[Fig. 2E, 2F]**. Consistent with previous studies, we observed age-dependent accumulation of VAT-T_R_ across all three genotypes in male mice (Feuerer et al., 2009), but despite considerable mouse-to-mouse variability, VAT-T_R_ remained elevated in YF and KO mice across all age groups analyzed **[Fig. 2G].**

**Figure 2:**
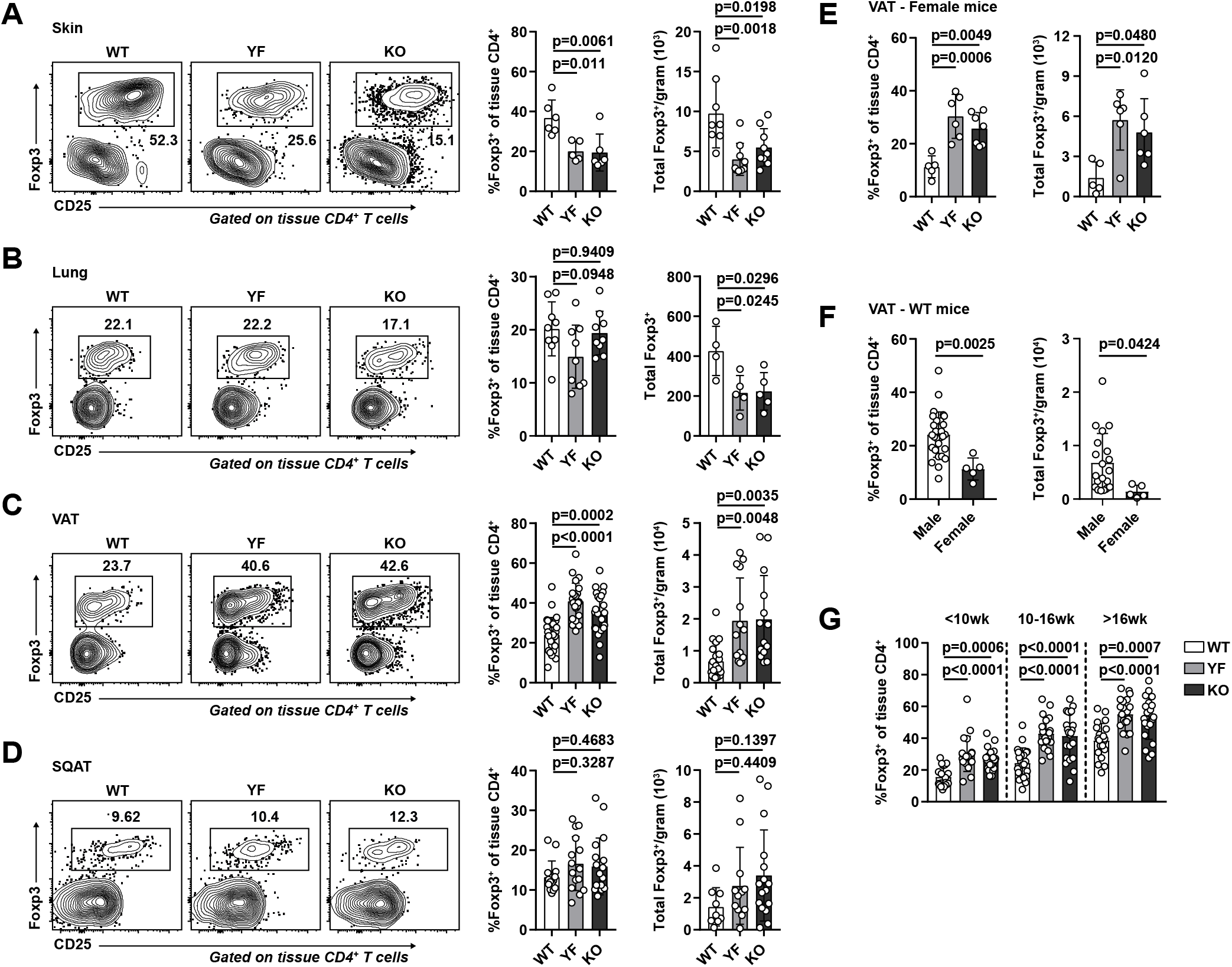
T_R_ frequency and number are increased in VAT in the absence of ICOS signaling. **(A-C)** Representative flow cytometry plots and quantification of of T_R_ among tissue-localized TCRβ^+^CD4^+^ cells in WT, YF, and KO ear skin **(A)**, lung **(B)**, VAT **(C)**, and inguinal SQAT **(D)** of male mice (Skin: *n* = 3 per group from 2 independent experiments. Lung: *n* = 3 per group from 3 independent experiments. VAT: *n* = 3-6 from 8 independent experiments. SQAT: *n* = 3-5 for 5 independent experiments). **(E)** Tissue-localized VAT-T_R_ frequency and total number in female WT, YF, and KO mice (*n* = 3 per group for 3 independent experiments). **(F)** T_R_ frequency and total number in VAT of male versus female WT mice (combined data from WT groups in **(C)** and **(E)**). **(G)** VAT-T_R_ frequencies by noted age bins in male mice as measured by flow cytometry (*n* = 3-5 mice per group from 18 independent experiments). Mice were age-matched within independent experiments and pooled data are from experiments using mice aged 8-16 weeks, except in **(G)** where age is indicated. Statistical significance was determined using one-way ANOVA with Tukey’s post-test or two-tailed Student’s *t* test where appropriate. All data are presented as mean values ± SD.

We utilized male mice for the remainder of our studies as we found female mice harbor ~4-5-fold fewer T_R_/gram VAT than age-matched males, and female mice are protected from the development of diet-induced metabolic syndrome (Ahnstedt et al., 2018; Pettersson et al., 2012; Vasanthakumar et al., 2020). The frequency of VAT-T_R_ expressing KLRG1 and CD69 was significantly elevated in YF and KO mice compared to WT controls **[Fig. 3A]**. By contrast, in the spleen there was a reduced frequency of YF and KO T_R_ expressed CD69, and very few T_R_ were KLRG1^+^ **[Fig. S2A]**. We assessed VAT-T_R_ Foxp3 and CD44 expression, and found no difference in the absence of ICOS **[Fig. S2B]**. We did however observe a modest increase in the expression of CTLA-4 and a slight reduction in the frequency of CD25^+^ T_R_ in YF and KO VAT **[Fig. 3A]**. Therefore, T_R_ in the VAT of YF and KO mice are enriched in expression of markers indicative of an eT_R_ and VAT-T_R_ phenotype (Feuerer et al., 2009; Smigiel et al., 2014b).

**Figure 3:**
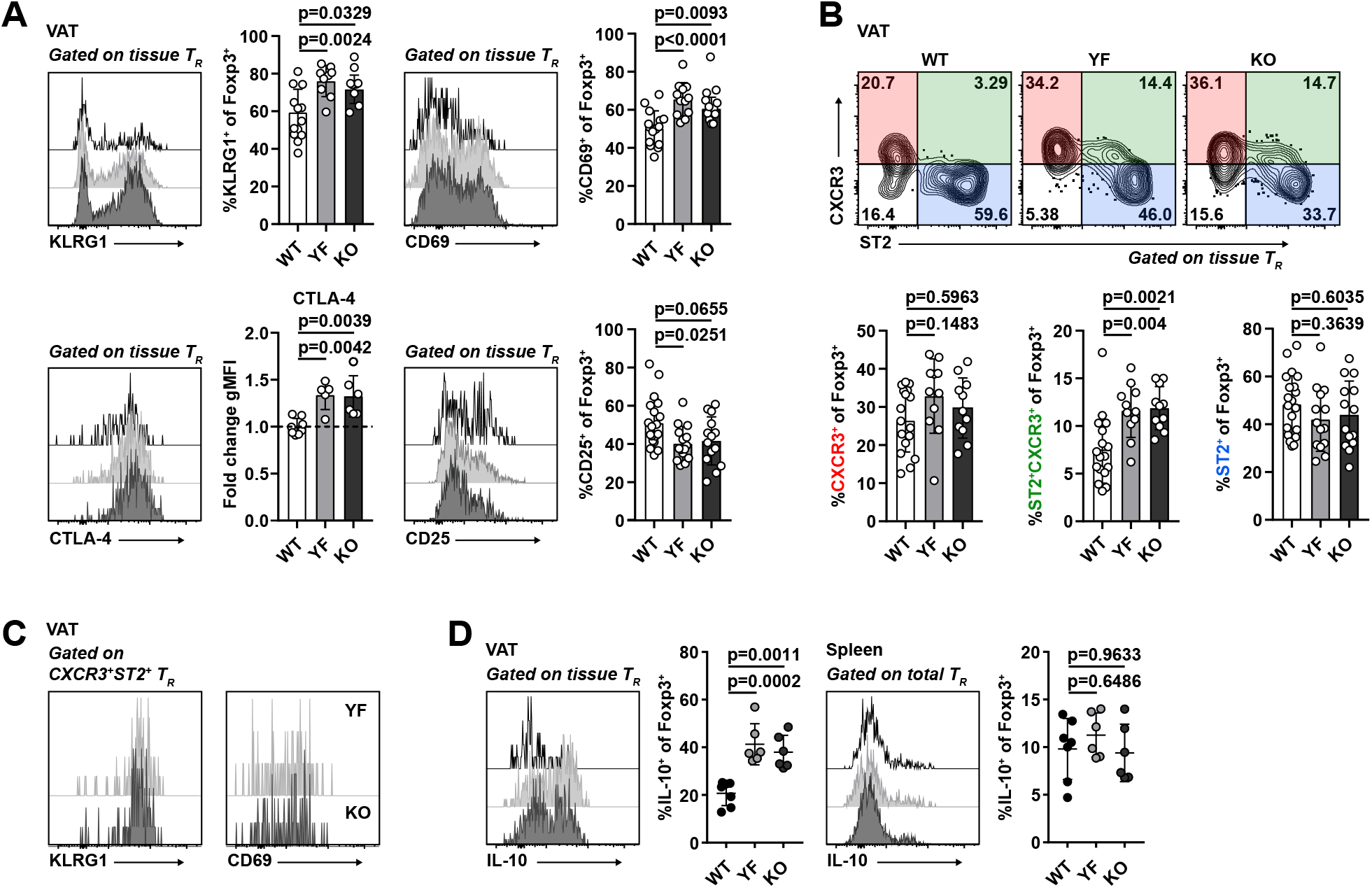
Loss of ICOS signaling supports an eT_R_ phenotype in VAT. **(A)** Representative flow cytometry plots and quantification of KLRG1, CD69, CTLA-4, and CD25 expression in tissue-localized VAT-T_R_. CTLA-4 gMFI was calculated as fold change compared to average expression in WT T_R_ for each individual experiment (*n* = 3-6 per group from 3 independent experiments or for CTLA-4, 2 independent experiments). **(B)** Representative gating of VAT-T_R_ on CXCR3 and ST2 expression and quantification of VAT-T_R_ positive for CXCR3 (left) or ST2 (right) alone, or double-expressers of CXCR3 and ST2 (middle) (*n* = 3-5 per group from 5 independent experiments). **(C)** Representative histograms showing expression of KLRG1 and CD69 in CXCR3^+^ST2^+^ YF and KO VAT-T_R_ as measured by flow cytometry. **(D)** Representative flow cytometry plots and quantification of IL-10 expression in T_R_ from VAT and spleen after 4h PMA/I+monensin stimulation *ex vivo* (*n* = 3-5 per group from 2 independent experiments). Mice were age-matched within independent experiments and pooled data are from experiments using male mice aged 8-16 weeks. Statistical significance was determined using one-way ANOVA with Tukey’s post-test. All data are presented as mean values ± SD.

In addition to prototypical eT_R_ markers, a fraction of VAT-T_R_ express the IL-33 receptor, ST2 (Kolodin et al., 2015; Vasanthakumar et al., 2015). Signaling downstream of ST2 is required for the establishment and maintenance of VAT-T_R_, and expansion of VAT-T_R_ through administration of IL-33 can reduce VAT inflammation and improve metabolic indices in models of diet-induced obesity (Han et al., 2015; Kolodin et al., 2015; Vasanthakumar et al., 2015). In addition to ST2-expressing T_R_, we found a substantial population of CXCR3^+^ T_R_ in the VAT of lean mice, confirming previously reported findings **[Fig. 3B]** (Li et al., 2018). Although there were no significant differences in the frequency of WT, YF, or KO VAT-T_R_ singly-expressing ST2 or CXCR3, we found a unique population of double-expressing ST2^+^CXCR3^+^ T_R_ in YF and KO VAT which resembled canonical VAT-T_R_ in their expression of KLRG1 and CD69 **[Fig. 3B, 3C]**. <u>Furthermore, whereas the abundance of splenic Treg cells in WT mice was dramatically reduced following treatment with the mTORC1 inhibitor rapamycin, there was no significant change in the numbe of Treg cells in the VAT **[Fig. S3]**, indicating that ICOS-dependent PI3K activation regulates VAT Treg cells independent of mTORC1 activity</u>.

ICOS signaling can potentiate IL-10 production (Busse et al., 2012; Dong et al., 2001; Herman et al., 2004; Hutloff et al., 1999; Kohyama et al., 2004; Kornete et al., 2012; Landuyt et al., 2019; Miyamoto et al., 2005; Redpath et al., 2013). Surprisingly, compared to VAT-T_R_ from WT mice, a substantially higher frequency of YF and KO VAT-T_R_ were poised to produce IL-10 upon *ex vivo* stimulation with PMA/I, with no appreciable differences in IL-10 expression by splenic T_R_ **[Fig. 3D]**. Thus, absence of ICOS-dependent PI3K signaling supports both the accumulation and suppressive phenotype of T_R_ in VAT.

### Baseline adipose inflammation is reduced in YF and KO mice

VAT inflammation is modulated through a balance of different immune cells. T_R_, eosinophils, anti-inflammatory adipose tissue macrophages (ATM), and group-2 innate lymphoid cells (ILC2) maintain a type-2 immune environment that supports metabolic homeostasis in lean animals. Inflammation however drives recruitment of neutrophils and cytotoxic T cells, differentiation of inflammatory ATM, and production of pro-inflammatory cytokines like TNFα (Burhans et al., 2018; Molofsky et al., 2013). Although there was no difference in mouse weights between WT, YF, and KO mouse across different ages, we did observe a trend toward increased VAT weight in YF mice, and a significant increase in KO mice **[Fig. 4A]**. By histology, we did not observe any overt morphological differences in VAT from WT, YF, and KO mice, including no observable differences in adipocyte size **[Fig. S4A].** The increase in VAT weight however did correlate with a modest increase in the total number of CD4^+^ T_eff_ per gram of tissue **[Fig. 4B]**. Despite this increase in VAT-T_eff_ number, there was a substantially lower frequency of YF and KO VAT-T_eff_ expressing features of inflammatory T_H_1 cells, including CXCR3, IFNγ, and TNFα **[Fig. 4C]**. Rather, ST2-expressing T_eff_ were enriched in YF and KO VAT **[Fig. 4C]**. Similar to VAT-T_R_, YF and KO VAT-T_eff_ were superior at producing IL-10 upon stimulation, suggesting that absence of ICOS signaling in effector cells supports a self-regulatory function **[Fig. 4C]**. IL-33 signaling induces production of T_H_2 cytokines (Schmitz et al., 2005), which maintain ATM in an anti-inflammatory state. While we didn’t find any significant changes in ATM or ILC2 populations **[Fig. S4B, 4C]**, we did note an increased frequency of eosinophils in YF and KO VAT **[Fig. 4D]**, which produce IL-4 to maintain anti-inflammatory ATM (Molofsky et al., 2013; Wu et al., 2011). Overall, global loss of ICOS expression and signaling impacts T_R_ along with other adipose-resident immune cells to maintain an anti-inflammatory state in VAT.

**Figure 4:**
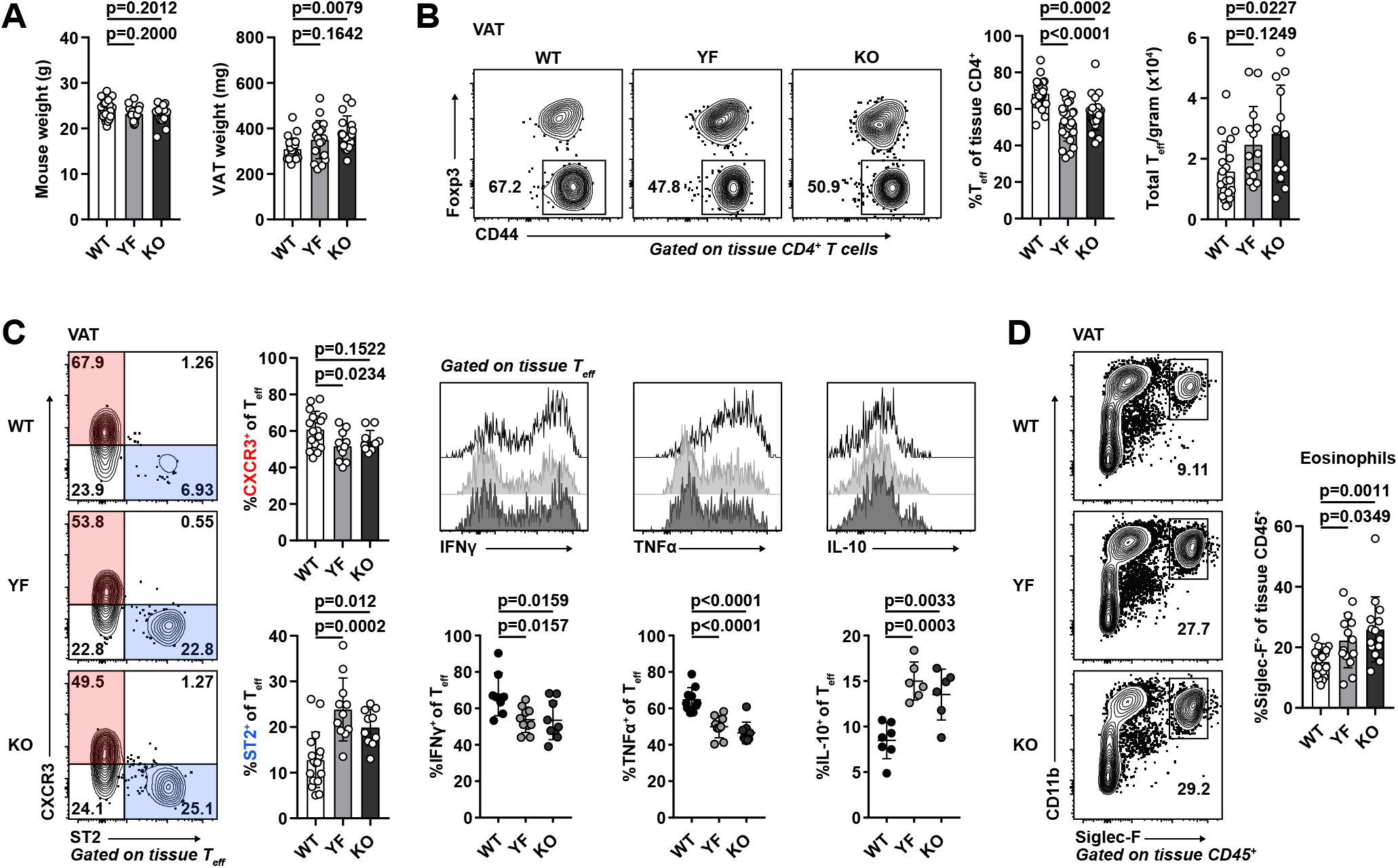
Reduced expression of inflammatory markers in YF and KO VAT. **(A)** Summary of total mouse and VAT weight in male WT, YF and KO mice (*n* = 3-6 per group from 6 independent experiments). **(B)** Representative flow cytometry plots and quantification of tissue-localized VAT-T_eff_ in WT, YF, and KO mice (*n* = 3-6 per group from 8 independent experiments). **(C)** Representative gating of VAT-T_eff_ on CXCR3 and ST2 expression and quantification of VAT-T_eff_ positive for CXCR3 or ST2 (left) (*n* = 3-5 per group from 5 independent experiments). Representative flow cytometry plots (top panels) and quantification (bottom panels) of IFNγ, TNFα, and IL-10 expression in VAT-T_eff_ after 4h PMA/I+monensin stimulation *ex vivo* (right) (*n* = 3-5 per group from 2 independent experiments). **(D)** Tissue-localized eosinophil frequencies in VAT (*n* = 2-6 from 6 independent experiments). Mice were age-matched within independent experiments and pooled data are from experiments using male mice aged 8-16 weeks. Statistical significance was determined using one-way ANOVA with Tukey’s post-test. All graphical data are presented as mean values ± SD.

### YF and KO mice are protected from high-fat diet-induced insulin resistance

Given the abundance and suppressive phenotype of T_R_ and other anti-inflammatory immune cells in VAT in YF and KO mice, we asked if they were more resistant to developing HFD-induced adipose inflammation and metabolic changes. Mice placed on an 18-week HFD steadily gained body mass, with a modest reduction in weight gained by KO mice compared to WT **[Fig. 5A]**. After 18 weeks on HFD, YF and KO mice sustained elevated T_R_ frequencies in VAT compared to WT mice, with an increased fraction of cells expressing the canonical VAT-T_R_ markers ST2 and KLRG1, and a smaller proportion of T_R_ expressing CXCR3 **[Fig. 5B, 5C]**. Additionally, YF and KO mice placed on HFD retained a greater frequency of eosinophils and T_eff_ skewed toward a type-2, ST2-expressing phenotype **[Fig. 5D, 5E]**. Fasting blood glucose levels (BGL) were similar among genotypes after 18-weeks, and WT, YF, and KO mice were able to clear glucose to the same extent, as measured by glucose tolerance test (GTT) **[Fig. 5F]**. However, in an insulin tolerance test (ITT), YF and KO mice demonstrated improved insulin sensitivity compared to WT mice **[Fig. 5G]**. Thus, loss of ICOS supports T_R_ and other anti-inflammatory cells in VAT during HFD, and this has functional consequences in maintaining metabolic homeostasis.

**Figure 5:**
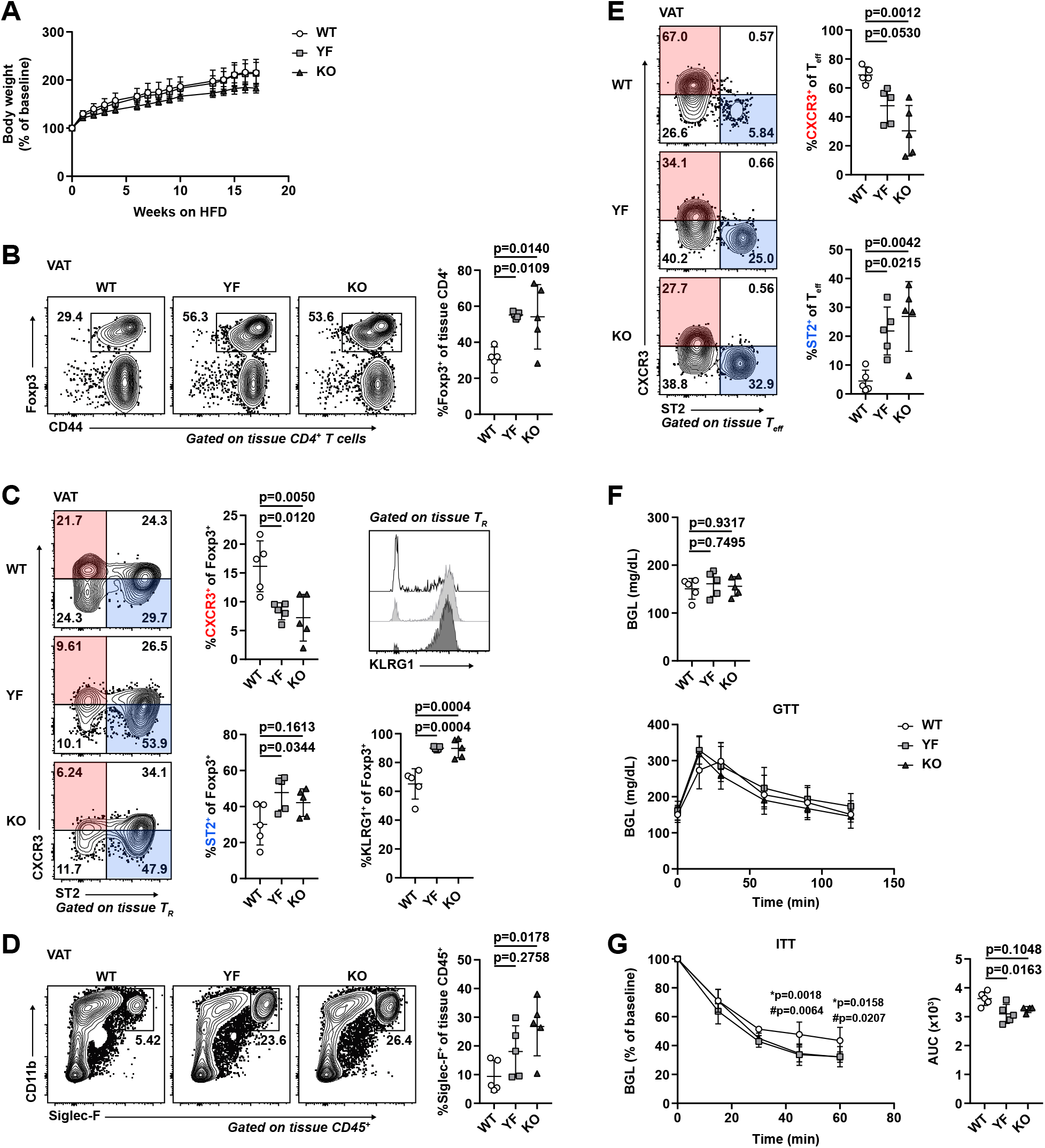
Mice deficient in ICOS signaling maintain anti-inflammatory immune cell abundance and phenotype in VAT after long-term HFD, correlating with improved insulin sensitivity. **(A)** Percentage of baseline body weight at indicated timepoints following initiation of HFD. **(B)** Tissue-localized VAT-T_R_ frequency after 18 weeks of HFD. **(C)** VAT-T_R_ phenotypic data as measured by flow cytometry showing expression of CXCR3, ST2, and KLRG1. **(D)** Frequency of tissue-localized eosinophils in VAT. **(E)** Representative gating of VAT-T_eff_ on CXCR3 and ST2 with summarized graphical data. **(F)** Fasting BGL (top) and GTT (bottom) in HFD-fed mice. **(G)** ITT in HFD-fed mice. * indicates significant difference between WT and YF; # indicates significant difference between WT and KO. ITT area under the curve (AUC) was calculated for each individual mouse. All data are representative of 2 independent experiments with *n* = 4-5 per group. For weight **(A)**, GTT, and ITT **(F, G)**, statistical significance was determined using two-way repeated measures ANOVA with Tukey’s post-test for multiple comparisons. For fasted BGL **(F)**, ITT AUC **(G)**, and all flow cytometry summary graphs, statistical significance was determined using one-way ANOVA with Tukey’s post-test. All graphs are presented as mean values ± SD.

### Absence of ICOS supports the accumulation and phenotype of VAT-T_R_ in a cell-intrinsic manner

ICOS signaling supports a wide array of cellular processes in the context of adaptive immunity (Panneton et al., 2019). To determine which of the T cell phenotypes we observed in intact YF and KO mice were the result of cell-intrinsic ICOS expression and signaling, we generated mixed bone marrow chimeras using congenically marked CD45.1^+^ WT and CD45.2^+^ WT, YF, or KO donors **[Fig. 6A]**. Body and adipose depot weights were similar between groups of chimeric mice **[Fig. S5A]**. Consistent with the increased abundance of VAT-T_R_ we observed at baseline in intact YF and KO mice, both CD45.2^+^ YF and KO T_R_ displayed a competitive advantage compared to CD45.1^+^ WT T_R_ in repopulating VAT in our chimeric mice. While WT:WT chimeras reconstituted VAT-T_R_ at a ~1:1 ratio, YF and KO T_R_ exhibited a significantly higher reconstitution ratio of ~5.5:1 **[Fig. 6B]**. In addition to supporting VAT-T_R_ abundance, absence of cell-intrinsic ICOS signaling also promoted a canonical VAT-T_R_ phenotype. Within the same chimeras, a higher frequency of YF and KO VAT-T_R_ expressed CD69 and ST2 **[Fig. 6C]**. Aligned with data from intact mice, more YF and KO VAT-T_R_ within chimeras were double-expressers of CXCR3 and ST2 **[Fig. 6C]**. Functionally, a larger proportion of YF and KO VAT-T_R_ were poised to produce IL-10 after *ex vivo* stimulation **[Fig. 6C]**. Mixed bone marrow chimeras retain a population of radioresistant host T_R_, which proliferate to replenish a portion of the available niche after irradiation (Komatsu and Hori, 2007), and this is particularly evident in many nonlymphoid tissues. Remarkably, in addition to preferentially reconstituting VAT compared to WT donor T_R_, YF and KO VAT-T_R_ were able to outcompete endogenous WT VAT-T_R_ as well, nearly replacing the entire T_R_ compartment in some chimeras **[Fig. 6D]**.

**Figure 6:**
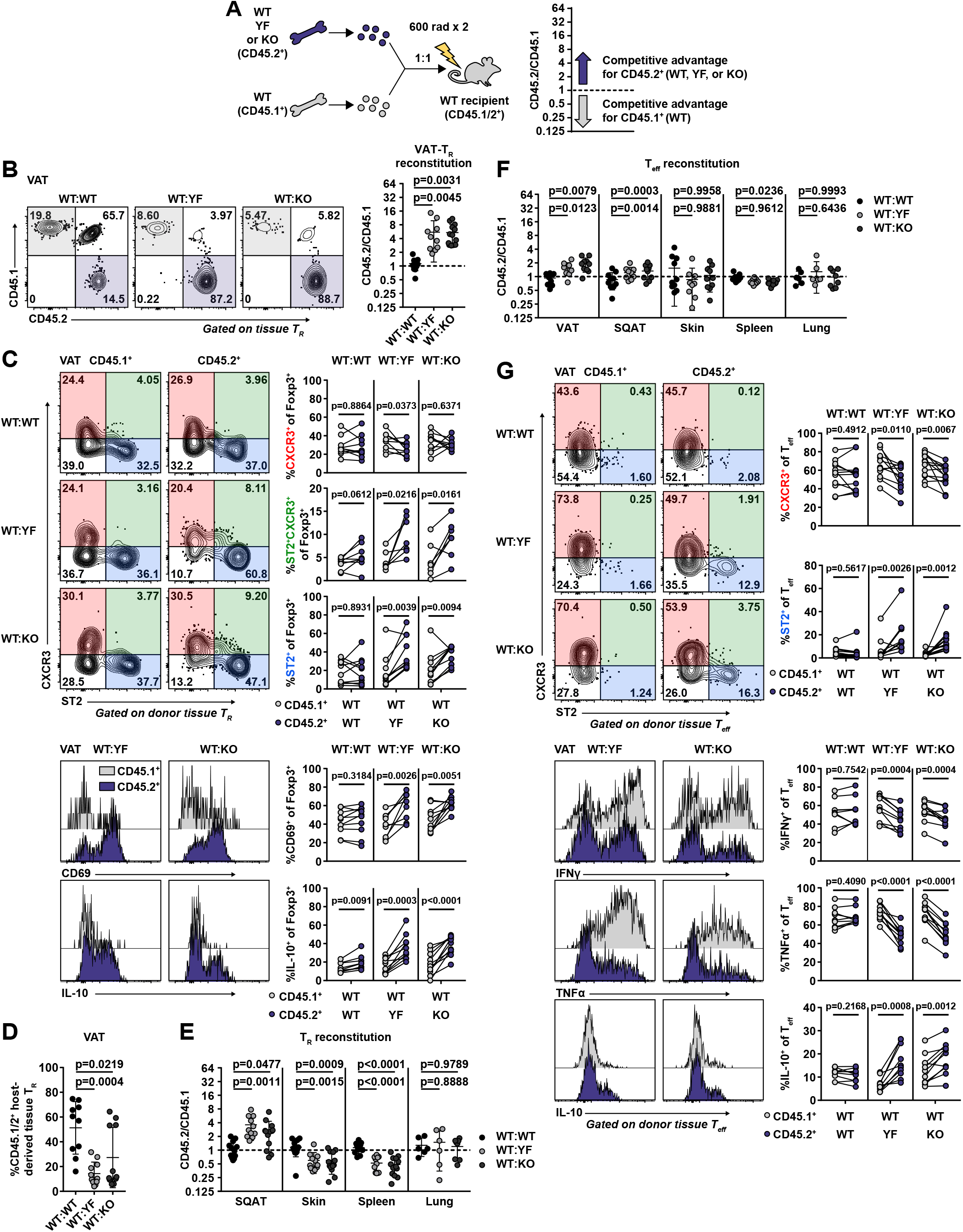
Cell-intrinsic ICOS signaling limits T_R_ and T_eff_ abundance and phenotype in VAT. **(A)** Schematic of mixed bone marrow chimera set-up (left) and analysis (right). **(B)** Representative gating of CD45.1 vs. CD45.2 expression in chimeric mice on tissue-localized T_R_ in VAT and quantification of donor T_R_ reconstitution in VAT. **(C)** Representative gating on CXCR3- and ST2-expressing CD45.1^+^ and CD45.2^+^ donor VAT-T_R_ in chimeric mice (top). Lines connect points indicating CD45.1^+^ and CD45.2^+^ cells within the same chimeric mouse. Histogram and quantification of CD69 expression in donor VAT-T_R_. Histogram and graphical summary of donor VAT-T_R_ expression of IL-10 after 4hr PMA/I+monensin stimulation *ex vivo* (bottom). **(D)** Frequency of endogenous CD45.1/2^+^ VAT-T_R_ in chimeric mice. **(E)** Reconstitution of donor T_R_ in the indicated tissues. **(F)** Summary of donor T_eff_ reconstitution in the indicated tissues. **(G)** CXCR3 and ST2 expression by gated VAT-T_eff_ (top). Histograms and summary of donor VAT-T_eff_ expression of IFNγ, TNFα, and IL-10 after 4hr PMA/I+monensin stimulation *ex vivo* (bottom). Data are pooled from 3 independent experiments with *n* = 3-5 per group. Statistical significance for cell population reconstitution was determined using one-way ANOVA with Tukey’s post-test for each tissue analyzed. Data are presented as mean values ± SD. A two-tailed, paired Student’s *t* test was used to assess statistical significance when comparing expression in donor cells within the same chimeric mouse.

Contrary to our findings in VAT, YF and KO T_R_ were at a selective disadvantage in the skin, spleen, and large intestine, with no difference in reconstitution in the lungs **[Fig. 6E, Fig. S5B]**. T_R_ found in SQAT are phenotypically distinct from VAT-T_R_ and do not appear to accumulate with age (Feuerer et al., 2009; Li et al., 2018). However, unlike in intact mice, we did note that YF and KO donor T_R_ displayed a competitive advantage in repopulating SQAT in chimeric mice at a ratio of ~3:1 **[Fig. 6E]**. This could be due to low level inflammation following irradiation of the recipient mice that drives reconstitution of the SQAT niche with T_R_ exhibiting a more VAT-T_R_ phenotype.

Although CD8^+^ T cells and ILC2s of either genotype were similarly represented **[Fig. S5C]**, we did find that YF and KO T_eff_ were modestly, but significantly, enriched in VAT, at a ~2:1 ratio compared to WT T_eff_ with no difference in reconstitution of other tissues **[Fig. 6F]**. A higher frequency of YF and KO VAT-T_eff_ were ST2^+^ within the same chimeras, with a reduction in CXCR3-expressing T_eff_ **[Fig. 6G]**. Functionally, YF and KO VAT-T_eff_ were comprised of fewer IFNγ^+^ and TNFα^+^ and more IL-10^+^ cells when stimulated *ex vivo* **[Fig. 6G]**, indicating that in addition to driving increased T_R_ abundance, the absence of ICOS signaling also promotes a regulatory phenotype in VAT-T_eff_. Taken together, loss of ICOS signaling supports VAT-T_R_, and to a lesser extent, T_eff_ accumulation and phenotype through cell-intrinsic mechanisms.

### ICOS signaling impacts expression of homing receptors in T_R_ that allow access to VAT

The increased abundance of VAT-TR from YF and KO mice could result from the combination of increased proliferation, increased survival, or increased migration to the VAT. The proliferation marker Ki67 was expressed by ~10-20% of VAT-T_R_, and we found no significant difference in the frequency of Ki67^+^ VAT-T_R_ between WT, YF, and KO mice **[Fig. 7A]**. Similarly, WT, YF, and KO VAT-T_R_ expressed similar levels of the pro-survival protein Bcl2 **[Fig. 7B]**. Despite the lack of differences in Ki67 or Bcl-2 expression, we cannot rule out that changes in cellular turnover contribute to the differences we see, as T_R_ express other pro-survival factors. However, recent data demonstrating continuous recruitment of T_R_ to VAT via the circulation led us to assess potential differences in migration of cells (Vasanthakumar et al., 2020).

**Figure 7:**
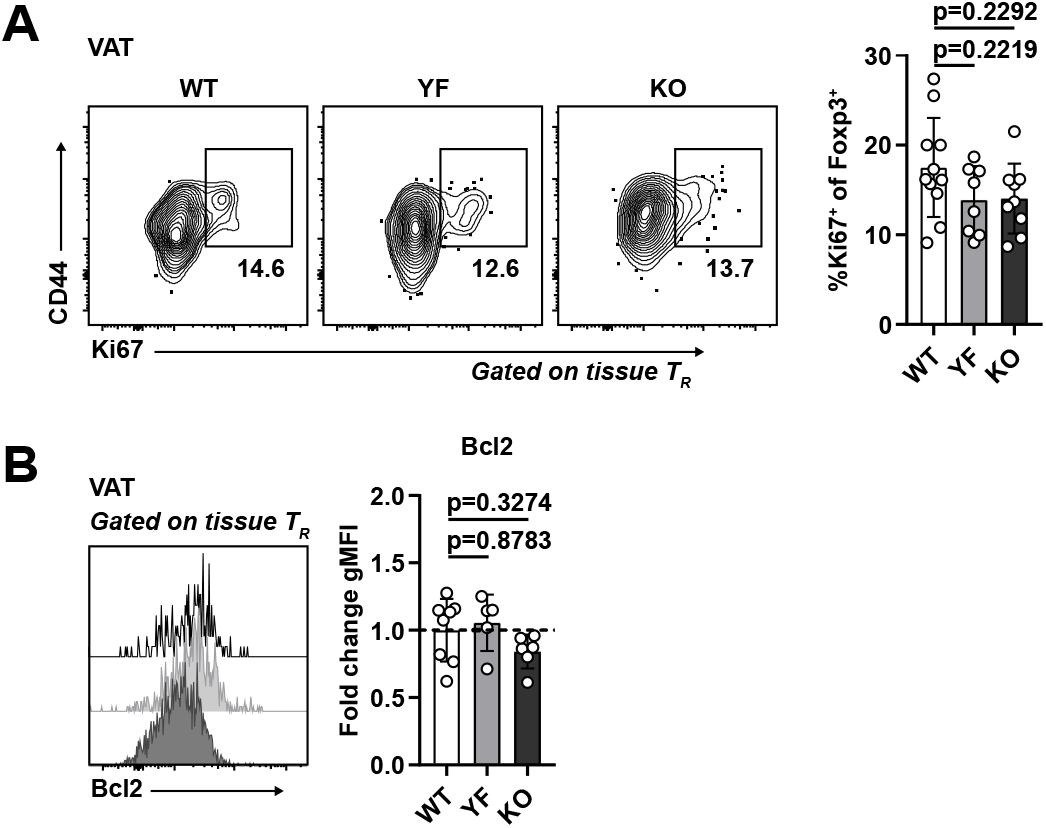
Increased abundance of YF and KO VAT-T_R_ is not due to enhanced proliferation or Bcl2 expression. **(A, B)** Ki67 **(A)** and Bcl2 expression **(B)** in tissue-localized VAT-T_R_ as measured by flow cytometry directly *ex vivo*. Bcl2 gMFI was calculated as fold change compared to average expression in WT T_R_ for each individual experiment (Ki67: *n* = 3-5 per group from 3 independent experiments. Bcl-2: *n* = 3-5 per group from 2 independent experiments). Mice were age-matched within independent experiments and pooled data are from experiments using male mice aged 8-16 weeks. Statistical significance was determined using one-way ANOVA with Tukey’s post-test. All data are presented as mean values ± SD.

In both intact and mixed bone marrow chimeric mice, YF and KO splenic T_R_ contained a small but significantly increased population of ST2^+^ T_R_ **[Fig. 8A, 8B]**, a subset of T_R_ that is both transcriptionally and epigenetically poised to take up residence in peripheral tissues (Delacher et al., 2017; Delacher et al., 2020). Given this, we hypothesized that superior migration and accumulation could account for the increased frequency of VAT-T_R_ in YF and KO mice. Although CCR2 facilitates recruitment of T_R_ to VAT (Vasanthakumar et al., 2020), we detected no difference in CCR2 expression among genotypes, and a modest decrease in CCR2^+^ T_R_ in KO spleens **[Fig. S6A]**, arguing against enhanced CCR2-mediated recruitment of YF and KO T_R_ to VAT.

**Figure 8:**
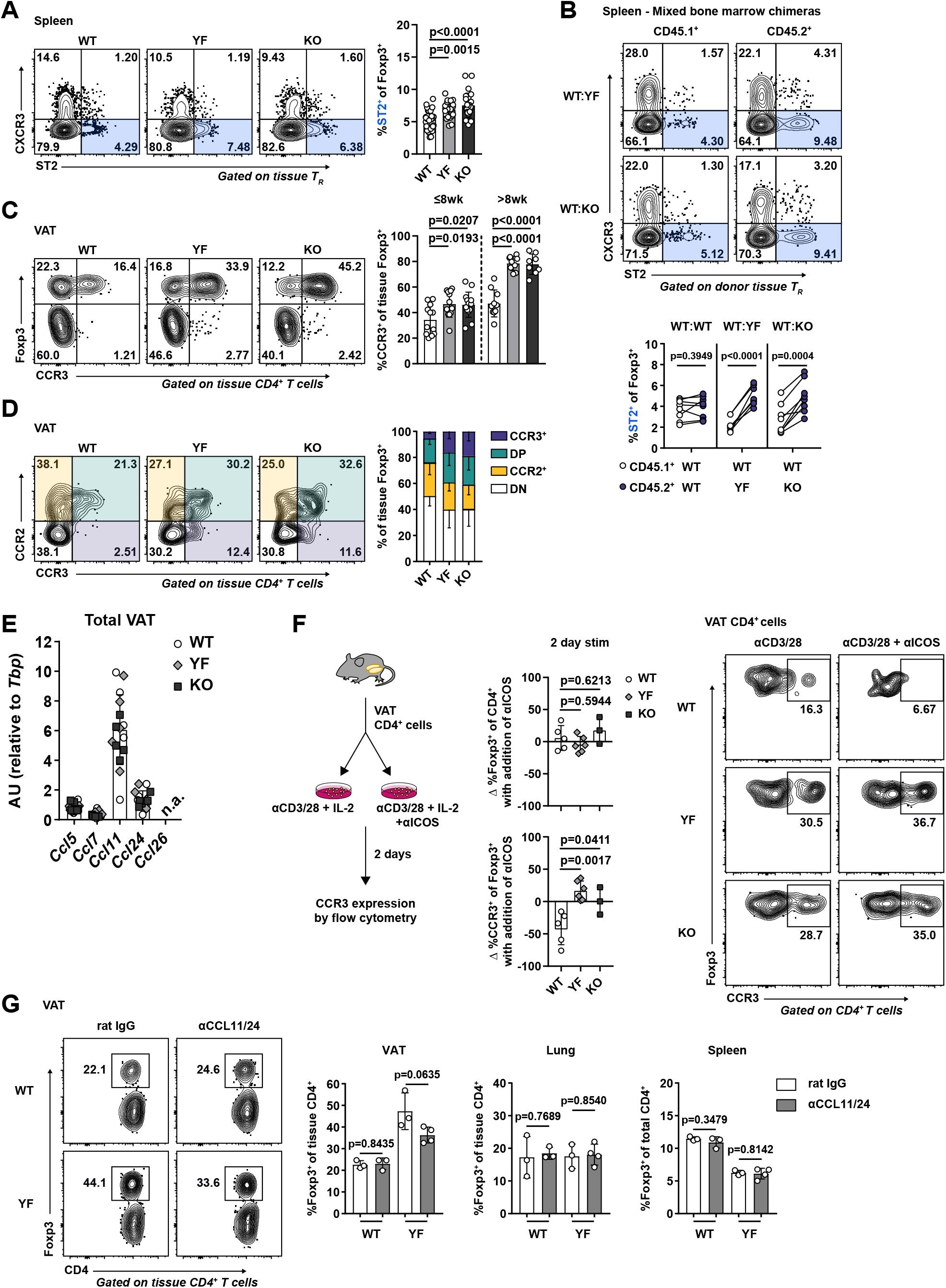
Increased VAT-T_R_ accumulation in the absence of ICOS signaling is associated with elevated CCR3 expression. **(A)** ST2 expression in splenic T_R_ (*n* = 3-5 per group from 8 independent experiments). **(B)** ST2 expression in CD45.1^+^ and CD45.2^+^ donor splenic T_R_ in WT:YF and WT:KO chimeric mice. Lines connect CD45.1^+^ and CD45.2^+^ donor splenic T_R_ within the same chimera (*n* = 3-5 chimeric mice per group from 3 independent experiments). **(C)** Expression of CCR3 in VAT-T_R_ in mice ≤8 wk and >8 wk of age (*n* = 3-5 per group from 7 independent experiments). **(D)** Expression of CCR2 and CCR3 by VAT-T_R_ (*n* = 3-5 per group from 2 independent experiments). **(E)** Expression of indicated CCR3 ligands in total VAT normalized to *Tbp* as measured by qPCR (*n* = 5 per group). **(F)** Schematic of *in vitro* culture experiments examining the impact of ICOS signaling on CCR3 expression (left). Graphs indicating fold change in T_R_ frequency of CD4^+^ cells and %CCR3^+^ of T_R_ between individual culture samples stimulated with or without αICOS for 2 d (middle). Representative flow cytometry plots with frequency of CCR3^+^ T_R_ after 2 d in specified culture conditions (right) (*n* = 1-3 per group from 3 independent experiments). **(G)**(Left) Representative flow cytometry plots indicating VAT-T_R_ frequency with or without CCL11/24 blockade. Graphs summarize T_R_ frequencies in indicated tissues after 2 wk (*n* = 3-4 per group). Mice were age-matched within independent experiments and collectively, pooled data are from experiments using male mice aged 8-16 weeks unless otherwise indicated. Statistical significance was determined using one-way ANOVA with Tukey’s post-test **(A, C)**, two-tailed, paired Student’s *t* test for expression in donor cells within the same chimeric mouse **(B)**, and two-tailed Student’s *t* test **(F, G)**. All data are presented as mean values ± SD.

Previous studies identified an open chromatin landscape around the *Ccr3* locus as well as high *Ccr3* transcript levels in VAT-T_R_ (Cipolletta et al., 2012; Li et al., 2018). The frequency of CCR3-expressing splenic T_R_ was slightly elevated in both young (≤8 weeks) and older (>8 weeks) YF and KO mice **[Fig. S6B]**, indicating the presence of a T_R_ population primed for both potential early seeding and continuous repopulation of peripheral tissues. In line with this, YF and KO VAT contained more CCR3^+^ T_R_ compared to WT mice and this population increased with age **[Fig. 8C]**. In mixed bone marrow chimeras, a significantly greater frequency of YF and KO donor T_R_ were CCR3^+^ in spleen and VAT **[Fig. S6C]**. Additionally, YF and KO VAT contained a unique population of T_R_ that expressed CCR3 alone, while WT VAT-T_R_ expressing CCR3 were also positive for CCR2 **[Fig. 8D]**. CCR3 expression was unique to VAT-T_R_, unlike CCR2, which we found to be also highly expressed on VAT-T_eff_ **[Fig. 8C, Fig. S6A]**. Additionally, the accumulation of CCR3^+^ T_R_ was unique to VAT, as we detected very little CCR3 expression in skin, lungs, and large intestine T_R_, and no differences across genotypes **[Fig. S6D, S6E].** We detected substantial expression of the CCR3 ligands *Ccl11* and *Ccl24* in the VAT, with no differences detected either by age or genotype and little expression of *Ccl5*, *Ccl7*, and *Ccl26* **[Fig. 8E]**.

To determine if ICOS signaling could directly impact expression of CCR3 on T_R_, WT, YF, and KO VAT-CD4^+^ cells were cultured with αCD3/αCD28 in the presence or absence of an agonistic αICOS antibody **[Fig. 8F]**. We additionally added 500 U/mL IL-2 to all conditions to account for any deficiencies in IL-2 production by YF and KO VAT-T_eff_. After 48 hr in culture, T_R_ frequency remained unchanged in both groups. However, the frequency of CCR3^+^ T_R_ in WT cultures significantly decreased after 2 days with addition of an ICOS agonist, whereas no changes were observed in YF or KO cultures, indicating that ICOS-dependent PI3K signaling can directly antagonize CCR3 expression in T_R_ **[Fig. 8F]**.

To address the impact of CCR3 signaling on continuous T_R_ seeding of VAT *in vivo*, we administered blocking antibodies to the CCR3 ligands CCL11 and CCL24 to 8-10 week old WT and YF mice. Blockade of CCL11/24 for ~2 weeks resulted in a modest although not statistically significant decrease in the frequency of VAT-T_R_ specifically in YF mice, with no differences in T_R_ abundance in spleen or lungs between treated and untreated mice **[Fig. 8G]**. We additionally noted a slight reduction in VAT-eosinophils in YF mice treated with αCCL11/24 **[Fig. S6F]**. Together, these results suggest a potential mechanism by which loss of ICOS-dependent PI3K signaling can drive increased recruitment of T_R_ to VAT by CCR3-CCL11/24 interactions.

## Discussion

We have identified surprising tissue-specific effects of ICOS signaling on T_R_ abundance and phenotype. Although there was a significant loss of CD44^+^ eT_R_ in the spleen, pLN, skin, and lungs of YF and KO mice compared to WT, there was a substantial increase in T_R_ abundance in VAT. T_R_ frequency and phenotype were similar in YF and KO mice, including comparably altered readouts of PI3K signaling, suggesting that ICOS primarily impacts T_R_ abundance and homeostasis via PI3K-dependent mechanisms.

In addition to supporting VAT-T_R_ abundance, loss of ICOS signaling promoted a highly suppressive VAT-T_R_ phenotype, including an increased proportion of T_R_ expressing KLRG1, CD69 and IL-10. *In vitro*, addition of ICOS stimulation augments IL-10 production in CD4^+^ T cells while loss of ICOS signaling *in vivo* results in reduced T_R_ abundance and IL-10 expression in models of autoimmunity, allergy, and infection (Busse et al., 2012; Hutloff et al., 1999; Kohyama et al., 2004; Kornete et al., 2012; Landuyt et al., 2019; Redpath et al., 2013). Although ICOS expression is associated with IL-10-producing T_R_, ICOS signaling does not appear to be intrinsically required for IL-10 production. CD4^+^ T cells from KO mice are capable of producing WT levels of IL-10 *in vitro*, and interestingly, appear to produce more IL-10 than WT counterparts under T_H_2-polarizing conditions (Dong et al., 2001). Furthermore, recent studies assessing T_R_ in the brain during chronic *Toxoplasma gondii* infection and lamina propria T_R_ at baseline report no changes in IL-10 production in the presence or absence of ICOS signaling (Landuyt et al., 2019; O’Brien et al., 2019). Therefore, the reduction in IL-10^+^ T_R_ with ICOS blockade observed in specific *in vivo* models is likely due to insufficient activation or access to specific environmental cues within tissues rather than an inherent requirement for ICOS signaling to drive IL-10 production.

VAT is a complex tissue, consisting of triglyceride-containing adipocytes and a variety of immune cells that function to maintain metabolic homeostasis. In YF and KO mice, other immune cells in addition to T_R_ showed evidence of an anti-inflammatory phenotype. Although ATM and VAT-ILC2 abundance was similar across genotypes at baseline, eosinophil frequency was significantly elevated in YF and KO VAT. In YF and KO mice, VAT-T_eff_ were skewed toward a T_H_2 phenotype in a cell-intrinsic manner, with an increased frequency of ST2-expressing and IL-10-producing cells. Therefore, in addition to promoting an anti-inflammatory state by sustaining VAT-T_R_, global loss of ICOS signaling supports the abundance and phenotype of other anti-inflammatory adipose-resident immune cells. These combined changes in cellular composition in VAT in the absence of ICOS signaling were maintained in the setting of long-term HFD, and rendered YF and KO mice more resistant to HFD-induced loss of insulin sensitivity. Changes in insulin responsiveness were subtle, despite a significant increase in T_R_ in YF and KO mice. A recent study demonstrated that ablation of IL-10 improved insulin sensitivity in mice (Rajbhandari et al., 2018). Thus it is possible that the increased IL-10 production we observe in VAT-T_R_ and VAT-T_eff_ in the absence of ICOS signaling offsets the insulin sensitizing effects of increased VAT-T_R_ abundance in these mice.

YF and KO donor T_R_ preferentially repopulated VAT in mixed bone marrow chimeras, and were able to outcompete both donor and endogenous WT T_R_. ST2 is expressed on a large fraction of VAT-T_R_ (Vasanthakumar et al., 2015), and although the frequency of ST2-expressing VAT-T_R_ was not changed in intact mice, YF and KO VAT-T_R_ in mixed bone marrow chimeras were comprised of significantly more ST2^+^ cells compared to WT donor T_R_, suggesting that the competitive advantage of YF and KO donor T_R_ in chimeric VAT is in part due to an enhanced ability to access available IL-33 through increased ST2 expression. The transcriptional regulators BATF and IRF4 cooperate to induce expression of ST2 in T_R_, possibly downstream of TCR signals as well as through IL-33 signaling itself (Vasanthakumar et al., 2015). Additionally, IL-2 can work synergistically with IL-33 to upregulate ST2 and specifically expand ST2^+^ T_R_ (Guo et al., 2009; Matta et al., 2014). T_R_ are unable to produce IL-2 themselves due to Foxp3-mediated transcriptional repression at the *Il2* locus (Wu et al., 2006; Ono et al., 2007) and therefore are dependent on paracrine sources of IL-2 produced by, for example, autoreactive T_eff_ (Setoguchi et al., 2005; Stolley and Campbell, 2016). However, KO T_eff_ are defective in production of IL-2 compared to T_eff_ from WT mice (Dong et al., 2001). The absence of an enriched population of ST2-expressing YF and KO VAT-T_R_ at baseline may be explained by this lack of IL-2, which is rescued in the presence of WT IL-2 producing cells in mixed bone marrow chimeras.

Molofsky *et al*. have argued that IL-33 signaling on T_R_ in VAT is mediated through ICOS/ICOSL interactions with ILC2s and that absence of this interaction results in loss of VAT-T_R_ (Molofsky et al., 2015). In their system, blockade of ICOS signaling was assessed in *Icosl*^*−/−*^ mice. Consistent with their data in *Icosl*^*−/−*^ mice, we did not observe any differences in ILC2 frequency or number in VAT of KO or YF mice. However, given that ILC2s express both ICOS and ICOSL, loss of one versus the other could differentially impact ILC2 function, resulting in unique cell extrinsic impacts on T_R_. Additionally, recent work revealed ligand-independent, constitutive ICOS signaling through PI3K-dependent pathways which relies on p85 association with the ICOS cytoplasmic YMFM motif and interactions between the ICOS transmembrane domain and the tyrosine kinase Lck (Feito et al., 2003; Wan et al., 2020). Thus the reported differences observed in our study and Molofsky’s may be the result of low level, cell intrinsic PI3K activation in ICOS-expressing T_R_ from *Icosl*^*−/−*^ mice.

Although CCR2 plays an important role in recruitment of T_R_ to VAT, we did not detect any differences in CCR2 expression in the absence of ICOS signaling. However, the frequency of CCR3^+^ T_R_ was increased in the spleen and VAT of YF and KO mice, which was due to cell-intrinsic loss of ICOS signaling and was enhanced with age. CCR3 is primarily associated with type-2 responses (Danilova et al., 2015; Francis et al., 2007; Humbles et al., 2002; Ma et al., 2002; Nagakubo et al., 2016), and lean, metabolically healthy VAT is maintained in an anti-inflammatory state by the presence of T_H_2-associated cells. *In vivo* blockade of CCL11/24 resulted in a modest reduction of T_R_ in YF VAT with no changes in other tissues analyzed. Parabiosis experiments demonstrate recruitment of circulating parabiont-derived T_R_ to VAT after months of shared circulation (Kolodin et al., 2015; Vasanthakumar et al., 2020), indicating that continual, low-level recruitment of T_R_ to VAT occurs throughout adulthood. Thus, longer-term antibody-mediated blockade may be necessary to observe significant effects on T_R_ accumulation in our system. Additionally, addressing the role of CCR3 is further complicated by its promiscuity, and whether compensation through other ligands occurs during *in vivo* blockade is unknown. *In vitro*, we found that ICOS stimulation resulted in reduced CCR3 expression in WT VAT-T_R_ with no changes in YF or KO cultures, suggesting that ICOS-dependent PI3K signaling inhibits expression of CCR3. Control of CCR3 expression in T cells has not been extensively characterized, however there is evidence that the transcription factor GATA3 is able to bind to a regulatory region in the *Ccr3* locus (Kong et al., 2013). Nonlymphoid T_R_, including VAT-T_R_, express high levels of GATA3 (Cipolletta et al., 2012; Delacher et al., 2020). However, whether ICOS signaling impacts CCR3 expression by modulating GATA3 expression or activity, and whether GATA3 is involved in controlling CCR3 expression in T_R_, remains to be explored.

Most studies assessing VAT-T_R_ biology utilize older male mice due to an increased abundance of canonical VAT-T_R_ with age (Cipolletta et al., 2015). However, mediators of tissue inflammation, including IL-6, TNFα, and IL1β, are upregulated in the VAT of aged male mice (older than 24 weeks) (Wu et al., 2007). We observed consistent accumulation of VAT-T_R_ from 5 weeks on in all three genotypes, which was correlated with an increase in CCR3^+^ T_R_ up to ~20 weeks of age. CCR2 regulates T cell access to sites of inflammation (Lee et al., 2007), and studies assessing the contribution of CCR2 on T_R_ migration to VAT utilize mice aged 25-30 weeks, when measures of VAT inflammation are increasing. CCR2 and CCR3 likely both have important roles for promoting recruitment of T_R_ to VAT, however it will be necessary to evaluate whether these receptors are utilized differentially based on age, sex, and inflammation. For example, CCR3 may be one important factor for early seeding and continuous recruitment of T_R_ to VAT in comparatively younger mice when VAT inflammation is low. With increasing age and inflammation (particularly in male mice), a switch to CCL2-CCR2 mediated migration may dominate.

Future studies are required to understand how ICOS exerts tissue-specific effects. The skin and lungs are under chronic stimulation by environmental antigens and commensal microbes, whereas the sterile VAT is maintained in a T_H_2-skewed environment in metabolically healthy mice. The milieu of chemokines and cytokines expressed at baseline in these tissues is likely very different. Indeed, T_R_ express distinct patterns of homing receptors and adhesion molecules to appropriately access specific nonlymphoid tissues (reviewed in Campbell & Koch, 2011). Absence of ICOS signaling may support expression of chemokine receptors that preferentially drive T_R_ to VAT due to constitutive expression of unique chemokines within this tissue. Additionally, VAT-T_R_ may rely less on PI3K signals compared to T_R_ found from other tissues, due to the presence of other supporting factors in the VAT of naïve mice, such as high levels of IL-33. Indeed, a recent study demonstrated that insulin receptor deletion in T_R_, which also results in reduced PI3K signaling, led to an increased suppressive phenotype in VAT-T_R_ and enhanced insulin sensitivity (Wu et al., 2020). <u>PI3K-mediated activation of mTORC1 also promotes glycolytic metabolism in T cells. Thus, the ability of the mTORC1 inhibitor rapamycin to diminish Treg cell abundance in the spleen but not the VAT suggests that Treg cells in different tissue sites may have distinct metabolic requirements for their homeostatic maintenance</u>.

We describe a surprising, antagonistic role for ICOS-dependent PI3K signaling in the accumulation, prototypical phenotype, and function of T_R_ specifically in VAT. We suggest that absence of ICOS signaling supports enhanced recruitment of T_R_ to VAT associated with increased expression of CCR3. Further studies will be needed to identify the molecular mechanisms by which ICOS inhibits the VAT-T_R_ phenotype and CCR3 expression. This highlights the complexity of tissue-specific T_R_ development and accumulation and the importance of considering how signals from the immune environment elicit both cell- and tissue-specific effects.

## Materials and Methods

### Mice

CD45.1^+^ B6.SJL (B6.SJL-*Ptprc*^*a*^*Pepc*^*b*^/BoyJ), B6.*Icos*^*−/−*^ (B6.129P2-*Icos*^*tm1Mak*^/J), and B6.*Foxp3*^*mRFP*^ (C57BL/6-*Foxp3*^*tm1Flv*^/J) mice were purchased from The Jackson Laboratory. B6.*Icos*^*Y181F*^ mice were provided by M. Pepper (University of Washington, Seattle, WA). All mice were backcrossed to a C57BL/6J background for at least 10 generations. *Foxp3*^*mRFP*^ mice were used as WT controls and mice from different genotypes used were co-housed at weaning. CD45.1^+^ B6.SJL mice were crossed to B6.*Foxp3*^*mRFP*^ mice to generate *Foxp3*^*mRFP*^ mice expressing CD45.1, CD45.2, and CD45.1/.2 allelic variants. B6.*Icos*^*−/−*^ and B6.*Icos*^*Y181F*^ mice were crossed to B6.*Foxp3*^*mRFP*^ mice to generate B6.*Icos*^*−/−*^ and B6.*Icos*^*Y181F*^ mice expressing *Foxp3*^*mRFP*^. All animals were bred and housed in specific pathogen-free conditions under the approval of the Institutional Animal Care and Use Committee of the Benaroya Research Institute._Males were used unless specified otherwise. Age of mice is indicated per experiment.

### Mixed bone marrow chimeras

Bone marrow cells were prepared from donor mice by flushing femurs and tibias with PBS. Red blood cells (RBC) from filtered cells were lysed in ACK lysis buffer (Life Technologies) for 5 min at room temperature (RT). Recipient mice were lethally irradiated (2 x 600 rad separated by ≥4 hr). Mixed bone marrow chimeras were generated by retro-orbitally (r.o.) injecting a 1:1 ratio (≥2 ×10^6^ million cells total) of bone marrow cells of the appropriate genotypes into anesthetized recipient mice. Chimeric mice were rested for ≥10 wk.

### Intravascular labeling

Mice were anesthetized with 4% isoflurane and 3 μg BV510- or BV650-conjugated CD45 (30-F11) was injected into mice r.o. in 200 μL PBS 5 min prior to sacrifice. Single cell suspensions were prepared for flow cytometry as described below, and localization of cells was determined by positive (blood-exposed) and negative (tissue-restricted) staining.

### Cell isolation

Single-cell suspensions were prepared from spleen and peripheral LNs (pooled inguinal, axillary, brachial, and superficial cervical nodes) by tissue disruption with frosted glass slides into RPMI with 10% bovine calf serum (R-10, BCS) and filtered through nitex mesh. Blood was collected via cardiac puncture into PBS containing 2 mM EDTA. Epididymal visceral adipose (VAT) and inguinal subcutaneous adipose (SQAT) were excised, finely minced with scissors, and digested in RPMI basal medium with 0.14 U/mL Liberase TM (Roche) and 10 U/mL DNase I (Roche) for 30 min at 37°C with shaking (200 rpm). Supernatants were filtered through a 100 μM cell strainers and washed several times to remove mature adipocytes from the stromal vascular fraction. Ears were harvested for skin-infiltrating cells. Dorsal and ventral sides were separated using forceps and digested as described for VAT/SQAT for 45 min. Lungs were digested as described for skin. Large intestines were cleaned, stripped of fatty tissue, and inverted. Tissue was placed into extraction media (RPMI basal medium, 2 mM DTT, 1 mM EDTA, 2% BCS) and shaken at 37°C for 20 min to release intraepithelial lymphocytes (IEL). IEL-containing supernatant was removed and filtered into 50 mL conical tubes. For lamina propria lymphocyte (LPL) isolation, immediately following incubation with extraction media, tissue was finely minced and placed into digestion media (RPMI basal media, 300 U/mL collagenase I [Worthington Biochemical Corporation], 1% BCS), shaken for 30 min at 37°C, and filtered into 50 mL conical tubes. Isolated cells from all tissues were incubated in ACK lysis buffer for 5 min, washed with R-10, and stained for flow cytometry or cultured for intracellular cytokine or chemokine receptor staining.

### Flow cytometry & intracellular cytokine staining

Single-cell suspensions were stained with Fixable Viability Dye eFluor 780 (eBioscience) in PBS for 10 min at RT. For surface staining, cells were incubated at 4°C for 30 min in FACS buffer (PBS, 2% BCS) with directly conjugated antibodies (Abs) for murine proteins. Abs purchased from BioLegend unless noted: CD4 (RM4-5), CD8 (53-6.7), CD11b (M1/70), CD11c (N418), CD25 (PC61), CD44 (IM7), CD45 (30-F11), CD45.1 (A20), CD45.2 (104), CD62L (MEL-14), CD69 (H1.2F3), CD206 (C068C2), CXCR3 (CXCR3-173), F4/80 (BM8), Gr1 (RB6-8C5), ICOS (15F9), KLRG1 (2F1/KLRG1), Siglec-F (E50-2440; BD Biosciences), ST2 (U29-93; BD Biosciences), TCRβ (H57-597). For intracellular staining, cells were stained for surface antigens as described above, washed, and permeabilized for 20 min with eBioscience Fix/Perm buffer at 4°C. Cells were washed and stained in PermWash buffer (eBioscience) for 30 min at 4°C with Abs (purchased from BioLegend unless noted) against Bcl2 (BCL/10C4), CTLA-4 (UC10-4B9), Foxp3 (FJK-16s; Invitrogen), IFNγ (XMG1.2; BD Biosciences), IL-10 (JES5-16E3), Ki67 (11F6), TNFα (MP6-XT22). For intracellular cytokine staining following restimulation, cells were incubated in FACS tubes with PMA (50 ng/mL) and ionomycin (1 μg/mL) plus monensin (2 μM) in 0.3 mL complete RPMI medium (RPMI-C) [RPMI with 10% (vol/vol) fetal bovine serum, 100 U/mL penicillin, 100 μg/mL streptomycin, 25 μg/mL gentamycin, 1 mM sodium pyruvate, 10 mM HEPES, 2 mM L-glutamine, 55 μM β-ME] for 4 hr at 37°C, 5% CO_2_ prior to staining as described above. Loss of mRFP expression occurred with our fixation/permeabilization protocols, requiring intracellular Foxp3 staining. Data were acquired on LSR II or FACSCanto flow cytometers (BD Biosciences) and analyzed using FlowJo software (TreeStar). Due to intraexperimental variability, geometric MFI (gMFI) was normalized as fold change compared to average expression in WT samples per experiment. Polybead polystyrene nonfluorescent microspheres (15 mm, Polysciences) were used to determine absolute cell numbers in flow cytometry samples. 100 μL of a fixed concentration (C_B_) of beads was mixed with 100 μL cells to be counted. Samples were acquired on a flow cytometer, with gates drawn on lymphocyte and bead populations based on their forward- and side-scatter properties. The ratio of lymphocyte gate events (N_L_) to bead gate events (N_B_) was determined and used to calculate the concentration (C_L_) of the original cell suspension as follows: C_L_ = (N_L_ / N_B_) x C_B_

### Phospho-flow cytometry staining

~1/4 of each spleen was immediately disrupted between frosted glass slides into BD Fix/Perm buffer (BD Biosciences). Cells were incubated for 20 min at RT, washed in FACS buffer, and resuspended in 90% ice cold methanol for ≥30 min. Cells were washed with BD Perm/Wash buffer (BD Biosciences) and stained for surface and intracellular antigens, including p-S6 (D57.2.2E; Cell Signaling Technology) for 45 min at RT.

### Chemokine receptor staining

Freshly isolated cells were incubated for 2.5 hr at 37°C, 5% CO_2_ in RPMI-C. For CCR7 expression, cells were incubated with CCL19-human IgGFc fusion protein for 20 min at 4°C, washed, then incubated with PE-conjugated goat anti-human IgGFc (Jackson ImmunoResearch Laboratories) for 20 min at 4°C. To detect CCR2 (475301; R&D Systems) and CCR3 (J073E5; BioLegend), cells were incubated for 30 min at 37°C with directly conjugated antibodies diluted in FACS buffer. Cells were washed twice with FACS buffer before being acquired on the flow cytometer.

### Histology

VAT was excised, immediately fixed in 10% formalin for 24 hr, then paraffin-embedded. Hematoxylin and eosin (H&E) staining was performed on 6 μm tissue sections by the Benaroya Research Institute Histology Core. Slides were imaged at room temperature with a Leica DM E brightfield microscope with a 10X eyepiece and 20X objective lens (together 200X). Images were caputured with a Leica EC3 3.1 megapixel digital color camera and imported into Leica Application Suite EZ imaging software.

### RNA extraction and quantitative PCR

VAT was dissected, immediately stabilized in RNAlater (Thermo Fisher), and frozen at −20°C until ready for processing. ~100 mg tissue was homogenized in 1 mL Qiazol and RNA was extracted using RNeasy Lipid Tissue Mini Kit (Qiagen) according to the manufacturer’s instructions. RNA quality and quantity was determined using an ND-1000 spectrophotometer (NanoDrop, Thermo Fisher). 500 ng template RNA was reverse transcribed with random hexamer primers in 20 μL using RevertAid First Strand cDNA Synthesis Kit (Thermo Fisher) and subsequently diluted 1:3.3 with nuclease-free water. qPCR was performed using 2 μL diluted cDNA and presynthesized TaqMan Gene Expression assays in TaqMan Gene Expression Master Mix (Applied Biosystems) for amplification of the following transcripts in a final volume of 20 μL: *Ccl5* (Mm01302427_m1), *Ccl7* (Mm00443113_m1), *Ccl11* (Mm0441238_m1), *Ccl24* (Mm00444701_m1), and *Ccl26* (Mm02763057_u1). Samples were run in technical triplicates using the QuantStudio 5 Real-Time PCR System (Thermo Fisher) with 10 min initial activation at 95°C followed by 40 cycles of 15 sec at 95°C, 60 sec at 60°C. Mean target mRNA levels were calculated by ΔΔCT method and normalized to Tbp (Mm01277042_m1) expression using QuantStudio Design and Analysis Software (Thermo Fisher).

### In vitro stimulation of VAT T cells

VAT was pooled from 2-3 mice of the same genotype to ensure adequate cell number for culture. Cells were isolated as described above, and CD4^+^ T cells were enriched by incubating cells with CD4 MicroBeads and positively selecting with MACS cell separation MS columns (Miltenyi Biotec). Purified cells were resuspended in RPMI-C with 500 U/mL recombinant IL-2 (eBioscience). Cells were cultured for 48 hr in 96-well flat-bottom plates with plate-bound αCD3 (2C11) and αCD28 (37.51) from Bio X Cell at 1 μg/μL, with or without the addition of plate-bound αICOS (C398.4A; BioLegend) at 2 μg/μL. Expression of CCR3 was assessed by flow cytometry as described above.

#### Rapamycin treatment

<u>Mice were given rapamycin (1mg/kg) three times per week for three weeks by intraperitoneal injection in PBS containing 5.2% PEG-400, 5.2% Tween 80, and 0.5% ethanol. Control animals were given vehicle only. Mice were euthanized for analysis of Treg cell abundance in the spleen and VAT 2 days after the final injection.</u>

### In vivo antibody blockade

Mice aged 8-10 weeks were given 0.75 μg/g body weight αCCL11 and αCCL24 (R&D Systems; MAB420 and MAB528) or an equivalent amount of rat IgG (Sigma) diluted in PBS by intraperitoneal (i.p.) injection on days 0, 5, and 10 and sacrificed for analysis on day 13.

### High-fat diet

Mice were placed on 18-week HFD (60% kcal fat diet, Research Diets D124928, *ad libitum*) beginning at 5-7 weeks of age. Weights and blood glucose (BGL) by tail prick (Contour Next One glucose meter; Ascensia Diabetes Care) were taken weekly after 6 hr of fasting.

### Insulin and glucose tolerance tests

Insulin tolerance tests were performed in conscious mice fasted for 6 hr. 1 U/kg human insulin (Sigma) was given i.p. and blood was collected by tail prick for BGL at 0, 15, 30, 45, and 60 min post insulin administration. Three days later, glucose tolerance tests were performed in conscious mice fasted for 6 hr. 1 g/kg glucose (Sigma) was administered i.p. Blood was collected by tail prick for BGL at 0, 15, 30, 60, 90, and 120 min post-glucose bolus.

### Statistical analysis

All data are presented as mean values ± SD and graphs were created and analyzed using Prism Software (GraphPad). Comparisons between genotypes were analyzed using one- or two-way ANOVA where appropriate, adjusted for multiple comparisons using Tukey’s post-test. For mixed bone marrow chimeras, statistical significance was determined using two-tailed paired Student’s *t* tests when comparing cells within the same chimeric mouse.

## Supplemental material

Fig. S1 shows ICOS expression across gated splenic T_R_ populations and summarizes pLN T_R_ frequency. Fig. S2 shows further characterization of splenic and VAT-T_R_ by flow cytometry. <u>Fig. S3 shows the impact of rapamycin treatment on Treg cell abundance in the spleen and VAT</u>. Fig. S4 shows VAT histology (H&E staining) and frequencies of ATM and ILC2 populations by flow cytometry. Fig. S5 shows further characterization of mixed bone marrow chimeras. Fig. S6 expands on CCR2 and CCR3 expression in specific tissues in both intact and chimeric mice (flow cytometry), shows expression of CCR3 ligands in whole VAT (qPCR), and characterizes eosinophil frequencies in VAT after *in vivo* CCL11/24 blockade

**Supplemental Figure 1 (goes with Figure 1): Despite normal surface expression of ICOS, YF mice phenocopy KO with equivalent loss of lymphoid T_R_. (A)** ICOS expression on gated WT, YF, and KO splenic T_R_. Fold change of ICOS gMFI is compared to average expression in WT T_R_ for each individual experiment (*n* = 3 per group from 5 independent experiments). **(B)** Representative ICOS expression on gated splenic KO and WT CD44^lo^CD62L^hi^ cT_R_ and CD44^hi^CD62L^lo^ eT_R_ as measured by flow cytometry (*n* = 3 per group from 5 independent experiments). **(C)** T_R_ frequencies of total CD4^+^ T cells in pLN (*n* = 2-5 per group from 5 independent experiments). Mice were age-matched within individual experiments and quantified data are pooled from experiments using male mice 8-16 weeks of age. Statistical significance was determined using one-way ANOVA with Tukey’s post-test. All data are presented as mean values ± SD.

**Supplemental Figure 2 (goes with Figure 3): Expression of activation markers on splenic and VAT-T**_**R**_. **(A)** KLRG1 and CD69 expression in gated WT, YF, and KO splenic T_R_ (*n* = 3-6 per group from 3 independent experiments). **(B)** CD44 and Foxp3 expression in tissue-localized VAT-T_R_. gMFI was calculated as fold change compared to average expression in WT T_R_ for each individual experiment (*n* = 3-6 per group from 6 independent experiments). Mice were age-matched within individual experiments and pooled data are from experiments using 8-16 week old male mice. Statistical significance was determined using one-way ANOVA with Tukey’s post-test. All data are presented as mean values ± SD.

**Supplemental Figure 3 (goes with Figure 3): Rapamycin treatment reduces Treg cell numbers in spleen but not VAT.** Number of CD4^+^Foxp3^+^ T reg cells in spleen (left) and VAT (right) in mice treated with rapamycin (1mg/kg) or vehicle as indicated (n=5 mice per group from one experiment). Statistical significance was determined using two-tailed unpaired Student’s *t* test. All data are presented as mean values ± SD.

**Supplemental Figure 4 (goes with Figure 4): Adipocyte, ATM, and ILC2 populations are unchanged in YF and KO VAT. (A)** H&E staining of VAT from 8 week old male mice. **(B)** Representative flow cytometry plots and quantification of tissue-localized ATM in WT, YF, and KO mice (top). Representative gating of ATM on CD11c and CD206 to identify M1-like and M2-like macrophages, respectively (bottom) (*n* = 2-6 per group from 6 independent experiments). **(C)** Frequency of tissue-localized ILC2s in VAT as measured by flow cytometry (*n* = 2-6 per group from 6 independent experiments). Mice were age-matched within individual experiments and pooled data are from experiments using 8-16 week old male mice. Statistical significance was determined using one-way ANOVA with Tukey’s post-test. All data are presented as mean values ± SD.

**Supplemental Figure 5 (goes with Figure 6): Cell- and tissue-specific changes in reconstitution in ICOS mutant mixed bone marrow chimeras. (A)** Summary of total mouse (left), VAT (middle), and SQAT (right) weights in mixed bone marrow chimeras. **(B)** Representative gating of CD45.1 vs. CD45.2 expression on tissue-restricted T_R_ in intraepithelial lymphocytes (IEL) from the large intestine. Graphs summarize reconstitution of donor T_R_ of IEL and lamina propria lymphocytes (LPL) in the LI. **(C)** Representative gating of CD45.1 vs. CD45.2 expression in chimeric mice on total CD8^+^ T cells (top) and ILC2s (bottom) in VAT. Graphs summarize reconstitution of donor CD8^+^ T cells (top) and ILC2s (bottom) in VAT. Data are pooled from 3 independent mixed bone marrow chimera experiments with *n* = 3-5 male chimeric mice per group. Statistical significance was determined using one-way ANOVA with Tukey’s post-test for each individual tissue. All data are presented as mean values ± SD.

**Supplemental Figure 6 (goes with Figure 8): Increased accumulation of CCR3**^**+**^ **T**_**R**_ **in the absence of ICOS signaling is specific to VAT. (A)** CCR2 expression by splenic (top) and VAT-T_R_ (bottom) as measured by flow cytometry (*n* = 3-5 per group from 2 independent experiments). **(B)** Expression of CCR3 in splenic T_R_ in mice ≤8 wk and >8 wk of age (*n* = 3-5 per group from 7 independent experiments). **(C)** Expression of CCR3 by gated CD45.1^+^ and CD45.2^+^ donor VAT-T_R_ in chimeric mice. Graphs summarize CCR3 expression by donor T_R_ in VAT and spleen. Line connects point representing CD45.1^+^ and CD45.2^+^ cells within the same chimeric mouse (*n* = 2-4 per group from 2 independent experiments). **(D)** CCR3 expression by tissue-localized skin T_R_ as measured by flow cytometry (*n* = 2-4 per group from 2 independent experiments). **(E)** CCR3 expression by CD45.1^+^ and CD45.2^+^ donor T_R_ within the same chimeric mouse in indicated tissues. Line connects CD45.1^+^ and CD45.2^+^ cells within the same chimeric mouse (*n* = 2-4 per group from 2 independent experiments). **(F)** Frequency of tissue-restricted VAT-eosinophils with and without *in vivo* CCL11/24 blockade as measured by flow cytometry (*n* = 3-4 mice per group). Mice were age-matched within individual experiments and pooled data are from experiments using 8-16 week old male mice. Statistical significance was determined using one-way ANOVA with Tukey’s post-test **(A, B, D)**, two-tailed, paired Student’s *t* test for expression in donor cells within the same chimeric mouse **(C, E)**, and two-tailed Student’s *t* test **(F)**. All data are presented as mean values ± SD.

## Acknowledgements

We thank A. Wojno, K. Arumuganathan, and T. Nguyen for maintaining the BRI Flow Cytometry Core; P. Johnson in the BRI Histology/Imaging Core; and D. Hackney and S. Zraika from the UW Diabetes Research Center Cell Function Analysis Core for help with metabolic profiling tests. M. Pepper generously provided *Icos*^*Y181F*^ mice with kind permission from W.-K. Suh. A. Burich, C. Toledano, and the BRI vivarium staff helped maintain mouse colonies. We thank members of the Campbell lab for helpful discussion and laboratory support.

This work was supported by grants to D.J.C. from the NIH (R01AI124693, R01AI136475). K.L.M. was supported by NIH NIAID T32 Grant AI106677, NIGMS T32 Grant GM007270, and the UW Diabetes Research Center Samuel and Althea Stroum Endowed Graduate Fellowship (2P30 DK17047).

## Author Contributions

K.L.M. and D.J.C. conceptualized study and designed experiments. K.L.M. performed experiments, analyzed data, and wrote original draft. D.J.C. reviewed and edited.

## Abbreviations

ATM: Adipose tissue macrophage
BGL: Blood glucose level
cT_R_: Central regulatory T cell
eT_R_: Effector regulatory T cell
HFD: High-fat diet
ILC2: Type 2 innate lymphoid cell
KO: *Icos*^*−/−*^
PI3K: Phosphoinositide 3-kinase
pLN: Peripheral lymph node
SQAT: Subcutaneous adipose tissue
T_eff_: Effector T cell
T_R_: Regulatory T cell
VAT: Visceral adipose tissue
YF: *Icos*^*Y181F*^

