## Supplementary figures and images for "ICOS signaling limits regulatory T cell accumulation and function in visceral adipose tissue"

### Supplemental Figure 1

#### Mittelsteadt *et al.* Supplemental Figure 1
#### (Goes with Figure 1)

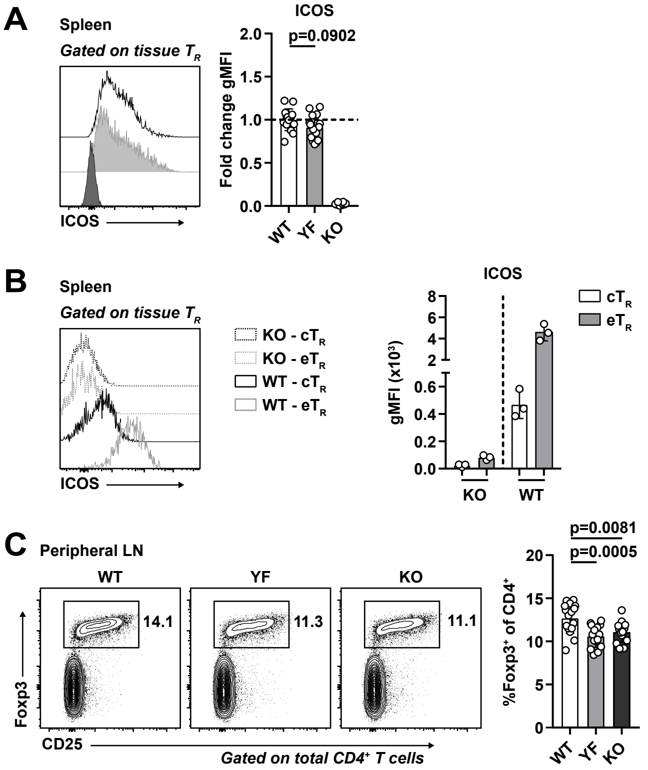

### Supplemental Figure 2

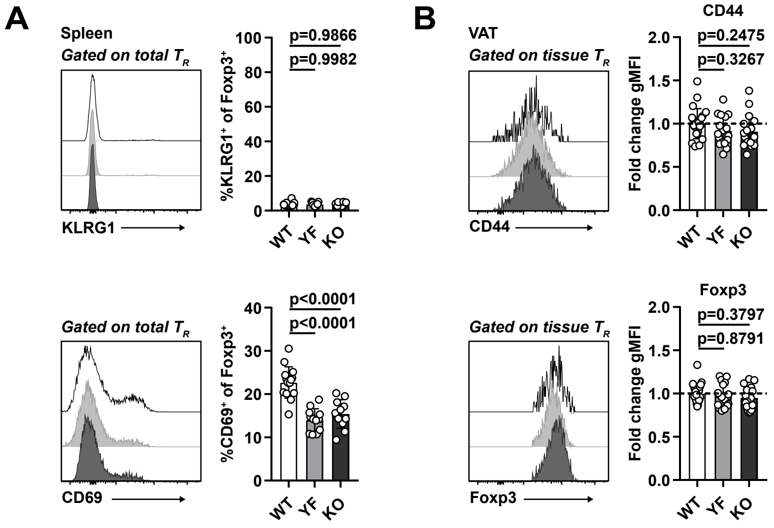

### Supplemental Figure 3

**Mittelsteadt et al. Supplemental Figure 3**
**(Goes with Figure 3)**

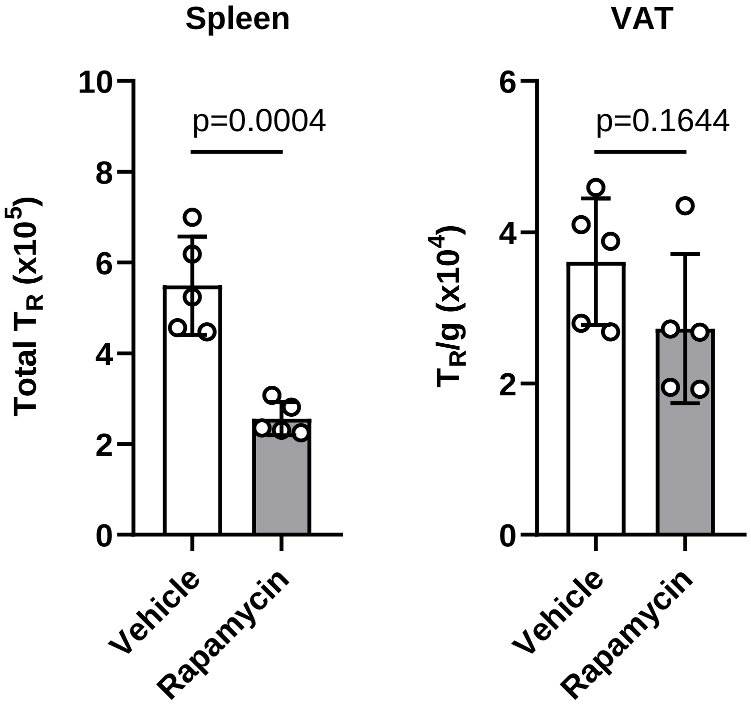

### Supplemental Figure 4

**Mittelsteadt *et al.* Supplemental Figure 4**
**(Goes with Figure 4)**

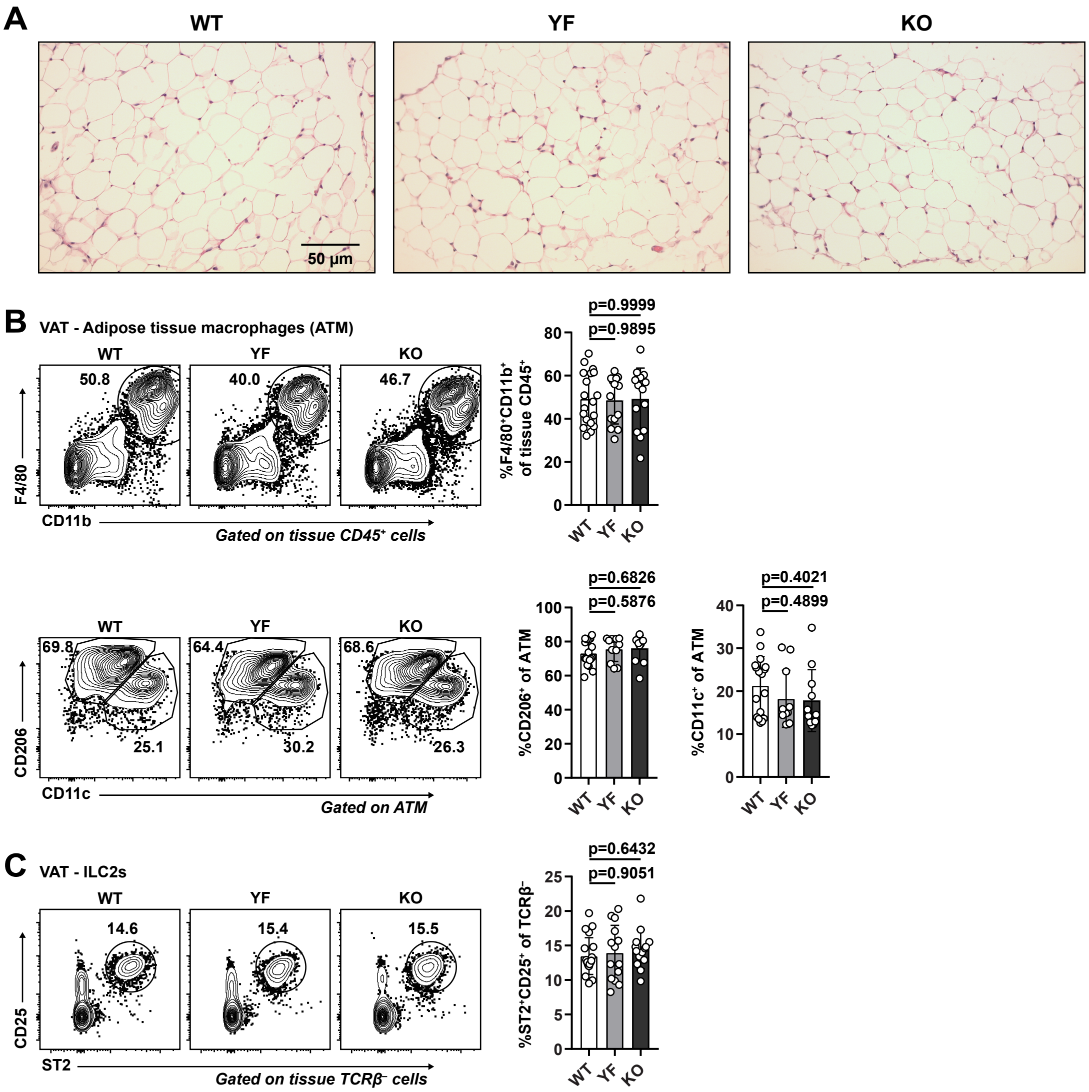

### Supplemental Figure 5

**Mittelsteadt *et al.* Supplemental Figure 5**
**(Goes with Figure 6)**

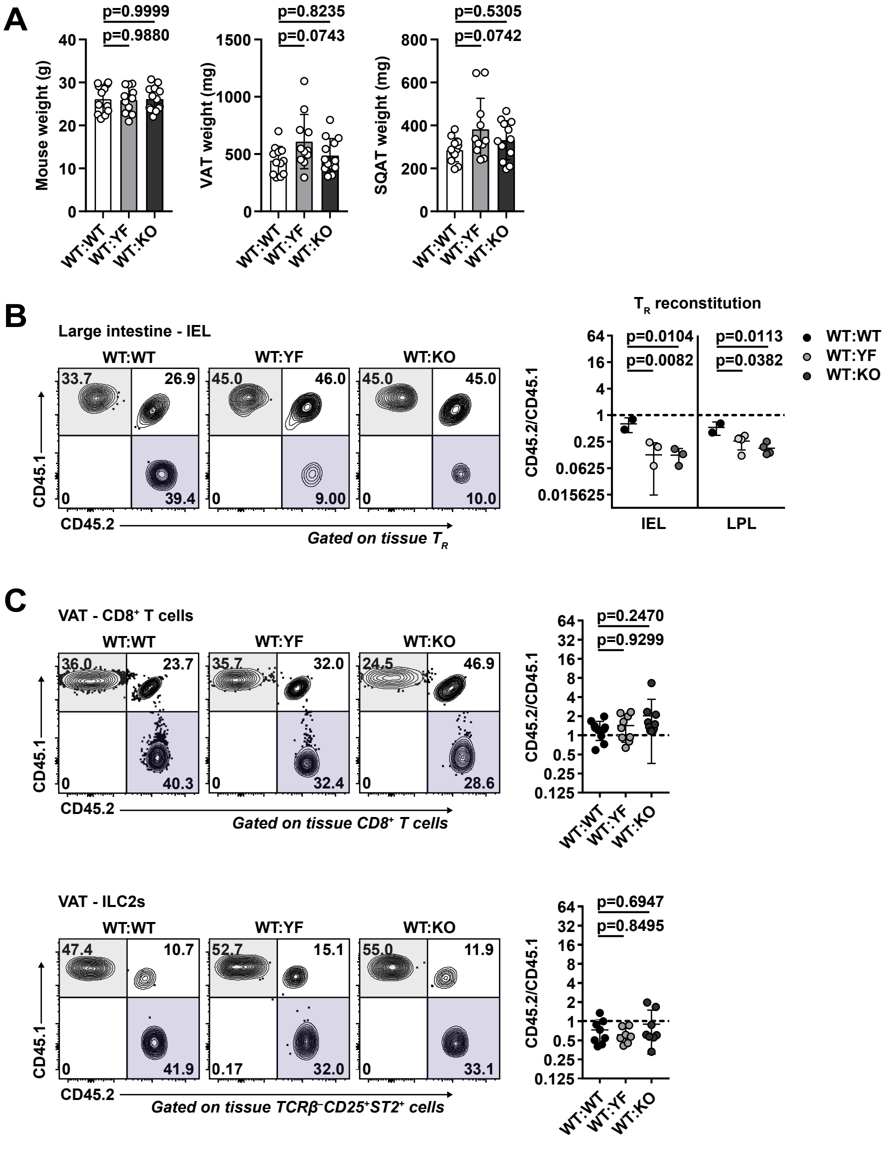

### Supplemental Figure 6

**Mittelsteadt *et al.* Supplemental Figure 6**
**(Goes with Figure 8)**

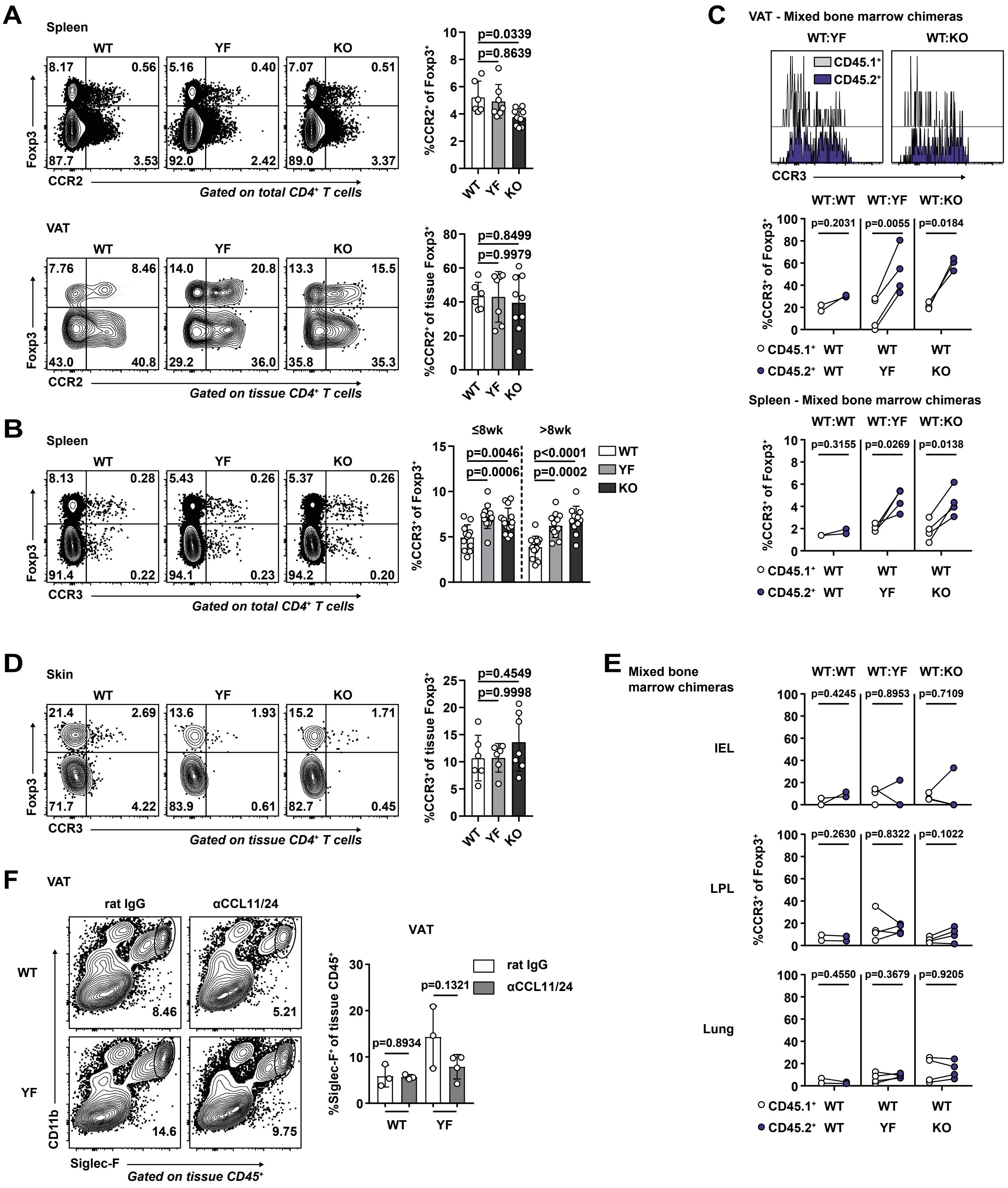
